# Sex-specific transcriptome of spinal microglia in neuropathic pain due to peripheral nerve injury

**DOI:** 10.1101/2021.05.27.446082

**Authors:** Nathan T Fiore, Zhuoran Yin, Dilansu Guneykaya, Christian D Gauthier, Jessica Hayes, Aaron D’Hary, Oleg Butovsky, Gila Moalem-Taylor

## Abstract

Recent studies have suggested a sexually dimorphic role of microglia in the maintenance of neuropathic pain in rodents. Here, we utilized RNA sequencing analysis and *in vitro* primary cultures of microglia to characterize the sex differences in microglia in pain-related regions in nerve injury and chemotherapy-induced peripheral neuropathy mouse models. Whilst mechanical allodynia and behavioral changes were observed in all models, transcriptomic analysis revealed a substantial change in microglial gene expression only within the ipsilateral lumbar spinal cord 7-days after nerve injury. Both sexes upregulated genes associated with inflammation, phagosome, and lysosome activation, though males revealed a prominent global transcriptional shift not observed in female mice, reflecting acute activation. Further, *in vitro* studies revealed that only male microglia from nerve-injured mice developed a reactive phenotype with increased phagocytotic activity. This study indicates distinct sex differences in spinal microglia and suggests they contribute to the sex-specific pain processing following nerve injury.

## INTRODUCTION

Chronic pain is a common public health issue that undermines the quality of life of chronic pain sufferers affecting 20% of the population above the age of 45, with a higher prevalence in women compared to men (Health and Welfare, 2020; Mogil, 2020). Neuropathic pain is a particularly debilitating form of chronic pain that comprises a wide range of heterogeneous conditions caused by nerve damage associated with traumatic injury, surgical intervention, various diseases, and anti-cancer treatments, such as chemotherapy (Finnerup et al., 2016; Kehlet et al., 2006). Despite various etiologies, neuropathic pain symptoms are similar in appearance and include sensory abnormalities (e.g. paresthesia and dysesthesia) and pain hypersensitivity (e.g. allodynia, hyperalgesia). Neuropathic pain is also characterized by behavioral disabilities (anhedonia, depression, exploration changes) that negatively impact the patient’s quality of life (Blyth et al., 2004; Mols et al., 2014).

A common feature in many pre-clinical models of neuropathic pain is neuroinflammation, and particularly glial activation (Austin and Moalem-Taylor, 2010; Lees et al., 2017). Recent evidence indicates the presence of sex differences in pain processing at all levels of the neuroaxis (Mogil, 2020), with a particular role of microglia (Sorge et al., 2015).

Microglia are central nervous system (CNS)-resident immune cells that constantly survey the microenvironment and maintain homeostasis. Microglia react to events (e.g. neuronal damage) that disrupt CNS homeostasis, which leads to rapid changes in their gene expressions and functional phenotypes. Recent genome-wide transcriptional studies have revealed a distinct molecular signature expressed in microglia during CNS homeostasis (Butovsky et al., 2014). This signature is lost during ageing, neurodegenerative diseases and inflammatory conditions, with a specific cellular phenotype appearing in a microglial subpopulation referred to as disease-associated microglia (DAM) or neurodegenerative microglia (MGnD), which is critical to the development of neurodegenerative conditions (Keren-Shaul et al., 2017; Krasemann et al., 2017; Sousa et al., 2018). Following damage to peripheral nerves, spinal and supraspinal microglia transition to reactive states in a time period correlated with sensory and affective-motivational pain behaviors, where they synthesise and release factors that facilitate neuronal excitability and transmission of nociception (Fiore and Austin, 2018; Gui et al., 2016; Tsuda et al., 2004). Whilst it is well accepted that microglia within the spinal cord and supraspinal pain-related regions have a critical role in the development of pain hypersensitivity (Gibson et al., 2019; Gui et al., 2016; Hu et al., 2018; Tsuda et al., 2004), recent studies have demonstrated sexually dimorphic microglial function in the development of neuropathic pain and pain relief in animal models (Doyle et al., 2017; Sorge et al., 2015). To further complicate the issue, most research over the years has only examined microglial changes in male mice (Mogil, 2020) and there are contradictory findings that observed no sex differences in the CNS after nerve injury (Lopes et al., 2017). Although previous reports showed sexual dimorphism in neuropathic pain, a transcriptome-wide assessment of gene expression in microglia in male and female mice has not been reported.

Here, we aimed to (i) establish whether there is a common microglia signature in different models of neuropathic pain; and (ii) determine if microglia display similar transcriptional responses between the sexes to further address the question of whether microglial sexual dimorphisms occur in the spinal cord or higher order supraspinal regions in neuropathic pain. To this end, we compared transcriptional changes of microglia from pain related CNS regions in male and female mice in the nerve injury model of sciatic nerve chronic constriction injury (CCI) and in two models of chemotherapy-induced peripheral neuropathy (CIPN) after paclitaxel and oxaliplatin treatments. By our approach, we have found no common microglia transcriptome in the different models of neuropathic pain and identified distinct sex differences in spinal microglia after CCI, which differ from other recently described microglial phenotypes.

## RESULTS

### Similar changes in pain behaviors in male and female mice after injury and chemotherapy

Mice were subjected to CCI of the sciatic nerve, paclitaxel- and oxaliplatin-induced peripheral neuropathy and were evaluated by various behavioral assays. All models of peripheral neuropathy exhibited neuropathic pain behaviors in male and female mice (**Figure 1**). Five days following CCI, 50% paw withdrawal threshold was significantly decreased in the ipsilateral (left) hindpaw compared to sham injured in both male and female mice (**Figure 1A**). Nerve-injured mice also showed behavioral changes as indicated by the open field holeboard test. There were no changes in distance travelled in male and female mice. However, male mice had reduced speed, nose pokes, rearing and time in the center of the field during the 5-minute task on day 6 after CCI compared to sham, whilst female mice had reduced speed, rearing and time spend in the center of the field **(Figure 2A)**. Following 6 cycles of Paclitaxel treatment, in both male and female mice, 50% paw withdrawal threshold was significantly decreased in the hindpaws compared to vehicle control mice on day 15 following initial injection (**Figure 1B**). Paclitaxel treated mice also showed behavioral changes as indicated by the open field holeboard **(Figure 2B)**. Both male and female mice had reduced time spent in the center of the field, whilst female mice also had reduced nose pokes 16 days after initial injection. Paclitaxel treated mice also had increased gait deficiency on day 15 following initial injection as indicated by an increase in the cumulative gait index (CGI) in both the forepaw and hindpaw after paclitaxel treatment **(Figure 1D)**. Following 12 cycles of Oxaliplatin treatment, in both male and female mice, 50% paw withdrawal threshold was significantly decreased in the hindpaws compared to vehicle control mice on day 22 after initial injection (**Figure 1C**). Oxaliplatin treated mice also showed behavioral changes as indicated by the open field holeboard test on day 23 after initial injection **(Figure 2C)**. Male and female mice had reduced distance covered, speed, rearing and nose pokes. Oxaliplatin treated mice also had increased gait deficiency as indicated by an increase in the CGI in both the forepaw and hindpaw in males and hindpaws in females on day 22 after initial injection (**Figure 1E**).

**Figure 1.**
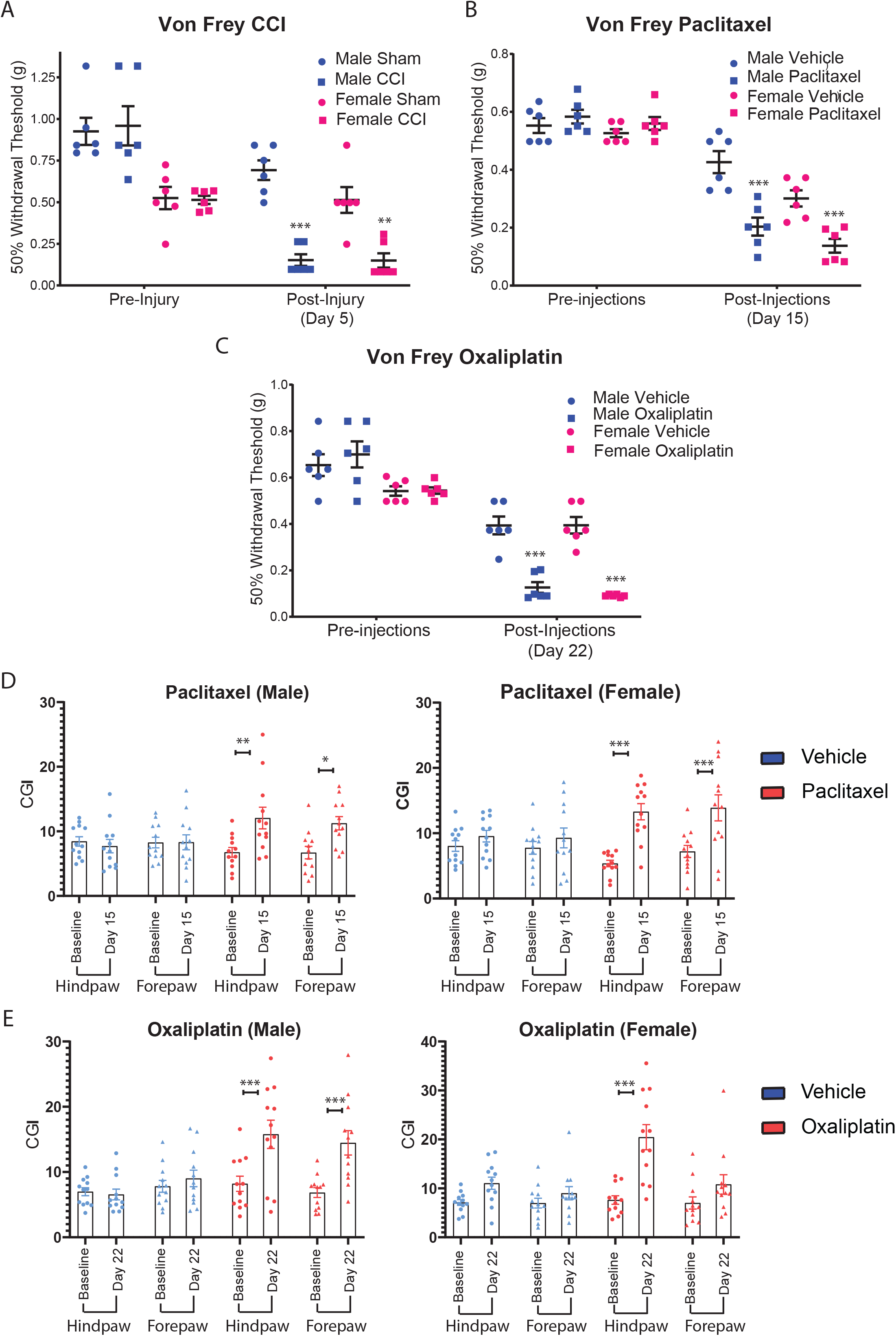
Comparable mechanical allodynia in males and females following CCI of the sciatic nerve and paclitaxel and oxaliplatin treatment. 50% mechanical withdrawal threshold was measured at baseline and (A) 5 days after CCI or sham surgery (left hindpaw), (B) 15 days after first paclitaxel injection or vehicle control and (C) 22 days after oxaliplatin treatment or vehicle control in both male and female mice using the up-down von Frey method (n=6 for each experimental group). Cumulative gait index (CGI) was measured using the DigiGait gait analysis system in vehicle control and (D) 15 days after paclitaxel treatment and (E) 22 days after oxaliplatin treatment (n=6 for experimental group). Data are shown as mean ± SEM (* p < 0.05, ** p < 0.01 and *** p < 0.001 for CCI or chemotherapy compared to sham or vehicle controls).

**Figure 2.**
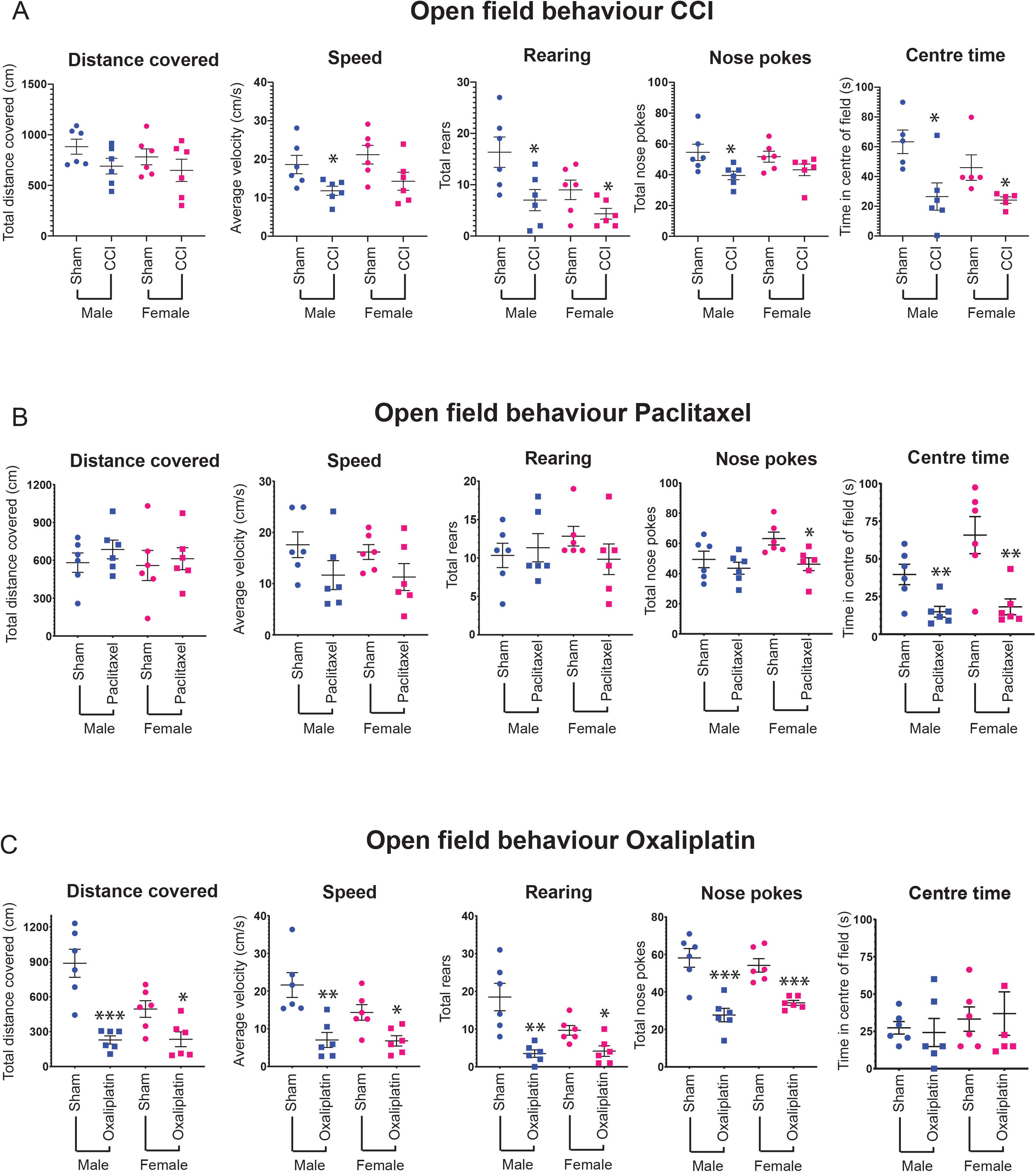
CCI of the sciatic nerve and paclitaxel and oxaliplatin treatments induce changes in locomotor activity and exploratory behaviors. The open field holeboard test was utilized to measure distance covered, average speed, number of rears and nose pokes as well as time spent in the center of the open field. Testing was conducted (A) 6 days after CCI, 16 days after paclitaxel treatment and (C) 23 days after oxaliplatin treatment. Data are shown as mean ± SEM (* p < 0.05, ** p < 0.01 and p < 0.001 for CCI or chemotherapy compared to sham or vehicle controls).

### Sex differences in spinal microglia transcriptome after CCI

Changes in microglia have been demonstrated in both the spinal cord and supraspinal regions in neuropathic pain (Austin and Fiore, 2019; Inoue and Tsuda, 2018). To investigate underlying common molecular mechanisms that regulate microglial dysfunction in neuropathic pain states in male and female mice, we isolated microglia from various CNS regions involved in pain processing one week after peripheral nerve injury or chemotherapy injection cycle, with a purity of >98% (**Supplementary Figure 1**). These regions included: (1) the ipsilateral lumbar spinal cord for CCI or all lumbar segment for CIPN (L3-L5); (2) the medial prefrontal cortex, hippocampus, and amygdala; (3) posterior thalamus and S1 cortex relating to the hindlimb; and (4) the periaqueductal gray and rostroventral medulla. We then analysed transcriptomes in male and female mice in CCI, paclitaxel and oxaliplatin models of peripheral neuropathy. We found high expression of known unique microglial genes (*P2ry12, Fcrls, Tmem119, Olfml3, Hexb* and *Tgfbr1*) in samples from all regions across our 3 models of neuropathy (data not shown). We analyzed differential gene expression (adjusted p < 0.05) in microglia between neuropathic and control mice across the different CNS regions. Using the Desq2 algorithm, we observed only few significant differences in differentially expressed genes (DEGs) in the oxaliplatin and paclitaxel models compared with vehicle controls. Similarly, after CCI only few significant changes were observed in DEGs in microglia within pain-related brain regions in male and female mice. **Table 1** summarizes these changes.

In contrast to supraspinal CCI regions and Oxaliplatin and Paclitaxel models, we identified 392 DEGs in combined male and female CCI ipsilateral spinal cord samples compared to male and female sham samples (adjusted p < 0.05), 309 of which were upregulated and 83 downregulated. Regarding sex-specific changes, transcription patterns in male and female spinal microglia after injury displayed some overlap. Male CCI mice had 210 DEGs compared to male sham mice (182 upregulated, 28 downregulated) and female CCI 96 DEGs compared to female sham mice (79 upregulated, 18 downregulated). The top genes (fold change > 2.5) are exemplified in **Figure 3A**. We validated these RNAseq results via quantitative real-time PCR (qPCR) of identified DEGs as well as identified microglial genes that do not change expression after CCI (**Supplementary Figure 2**). Of the DEGs observed in CCI ipsilateral lumbar spinal cord samples, 45 DEGs are common amongst male and female CCI mice, 165 DEGs are exclusive to male CCI mice (79%) and 51 are exclusive to female CCI mice (53%). Specifically, we found that 44 upregulated DEGs were common between male and female CCI mice, 138 are exclusive to male (76%) and 35 are exclusive to female mice (44%). Regarding downregulated genes, only 1 gene is common amongst male and female mice after CCI, 27 DEGs are exclusive to male (97%) and 16 DEGs are exclusive to female mice (94%) (**Figure 3B**). K-means clustering was used to identify DEGs with varying expressions between male and female mice compared to their controls. This analysis revealed 10 expression patterns, demonstrating a clear sexual dimorphism in response to nerve injury in the ipsilateral spinal cord (**Figure 3C**). Clusters with notable patterns include; Cluster 1: CCI male and female upregulated, though male upregulation more prominent; Cluster 2: CCI male and female upregulated, though female more prominent; Cluster 4: CCI male upregulated and CCI female no change or downregulated; Cluster 6: CCI male and female downregulated, though male more prominent; Cluster 7: CCI male and female downregulated, though female more prominent and; Cluster 9: CCI male downregulated and CCI female no change or upregulated. DEGs from these notable clusters are included in **Supplementary Table 1**. To identify the biological processes and molecular pathways differentially regulated in males and females after CCI, we subjected the DEGs (adjusted p value < 0.05) to Gene Ontology (GO) and Kyoto Encyclopedia of Genes and Genomes (KEGG) Pathway enrichment analysis using DAVID. Sexually dimorphic regulation of biological processes, with some overlap, was detected. GO enrichment analysis of upregulated genes in spinal microglia common in male and female mice revealed significant involvement (adjusted p < 0.05 and gene count > 5) in *immune system process* and the *innate immune system* (**Figure 3D**). The enriched KEGG pathways in upregulated common DEGs were *Lysosome, Phagosome, Antigen processing and presentation, Staphylococcus aureus infection and Tuberculosis* (**Figure 3E**). There was no significant enrichment of downregulated genes identified. Regarding the exclusively male CCI upregulated DEGs, GO identified significant involvement in *Translation, Cytoplasmic translation, rRNA processing, Immune system process, Ribosomal small unit assembly, Innate immune system* and *ribosomal small unit biogenesis* (**Figure 3D**) and KEGG pathway association with *Ribosome* (**Figure 3E**). On the other hand, GO of the corresponding female exclusive DEGs uncovered significant involvement in *Transport* and no association with KEGG pathways (**Figure 3D, E**).

**Figure 3.**
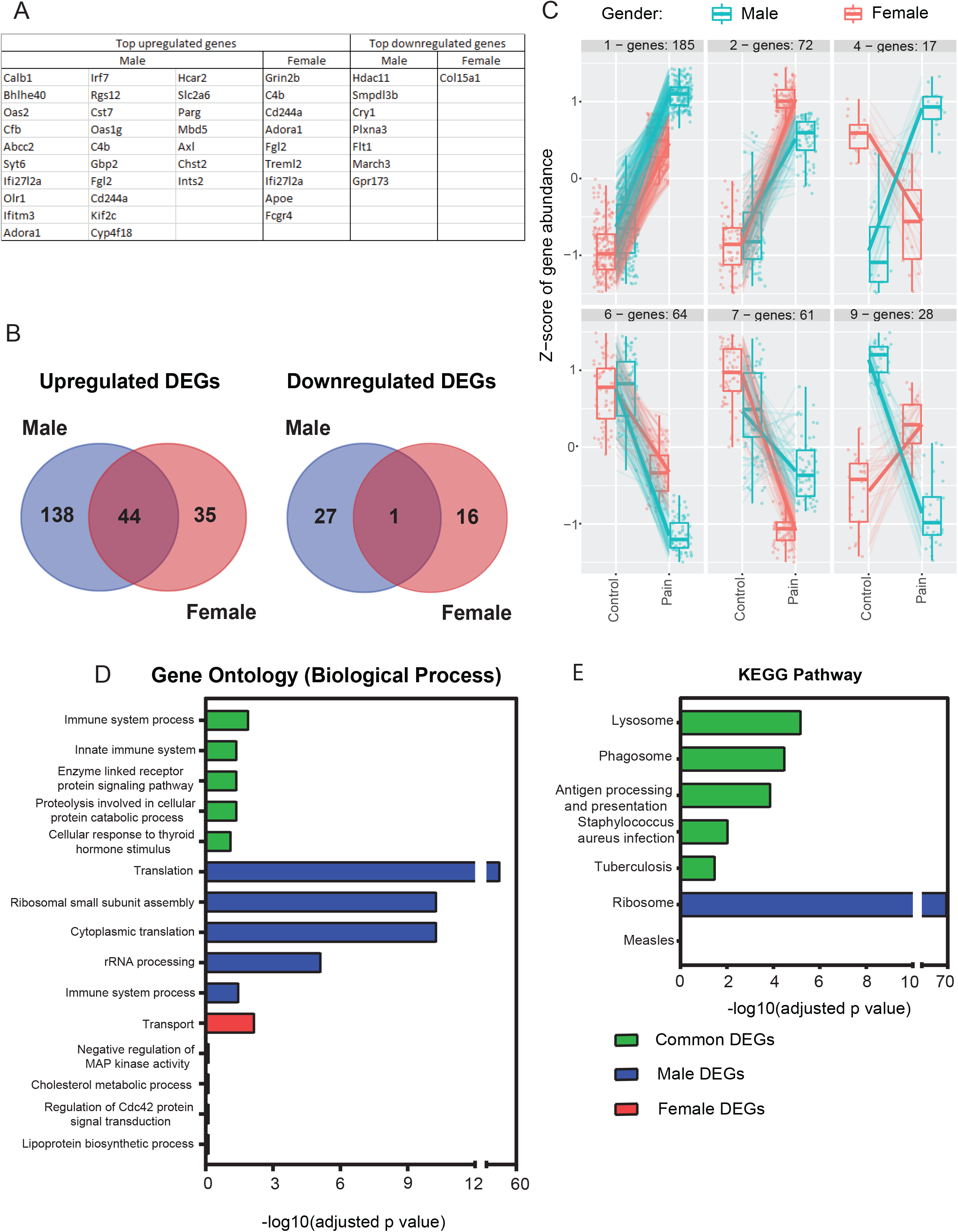
Sex differences in spinal microglia after CCI. (A) A list of the top differentially expressed genes (ranked by fold change) in male and female mice after CCI. (B) Venn diagrams highlighting common and exclusive upregulated and downregulated DEGs found in male (blue) and female (red) microglia in the ipsilateral lumbar spinal cord 7 days after CCI. Genes were clustered (k means = 10) based on variance in male and female z score differences after CCI. The top (D) GO enriched biological functions and (E) KEGG pathways from DEGs that are upregulated in both male and female mice (Common DEGs), only upregulated in males (Male DEGs) or only in females (Female DEGs) 7 days after CCI in spinal microglia. Data are shown as Benjamini-Hochberg adjusted p value.

To better understand the physiological function of the transcriptomic changes observed in male and female spinal microglia following CCI, the ipsilateral spinal microglia RNA-seq data were submitted to Ingenuity Pathway analysis (IPA) core analysis (DEGs with a fold change > [1.5] and adjusted p < 0.2 were included). These differentially expressed genes were categorized to related upstream regulators (**Supplementary Table 2** and **3**), canonical pathways (**Supplementary Table 4** and **5**), diseases and functions (**Supplementary Table 6** and **7**). Notable upstream regulators of male spinal microglia after CCI include activation of IRF3 (interferon regulatory factor 3) and IRF7, and suppression of SOCS1 (suppressor of cytokine signaling 1), whereas female microglia include activation of TREM2 (triggering receptor expressed on myeloid cells 2) and IFNG (interferon-γ) and suppression of IL10RA (Interleukin 10 Receptor Subunit Alpha) (**Figure 4A**). Notable changes in canonical pathways in male CCI microglia include activation of EIF2 (eukaryotic initiation factor 2) signaling and interferon signaling and inhibition of TGF-β (transforming growth factor beta) signaling (**Figure 4B**), whilst only a few canonical pathways were significantly altered in female microglia (**Figure 4B**). The top significantly altered diseases and functions for male and female microglia are summarized in **Figure 4C**.

**Figure 4.**
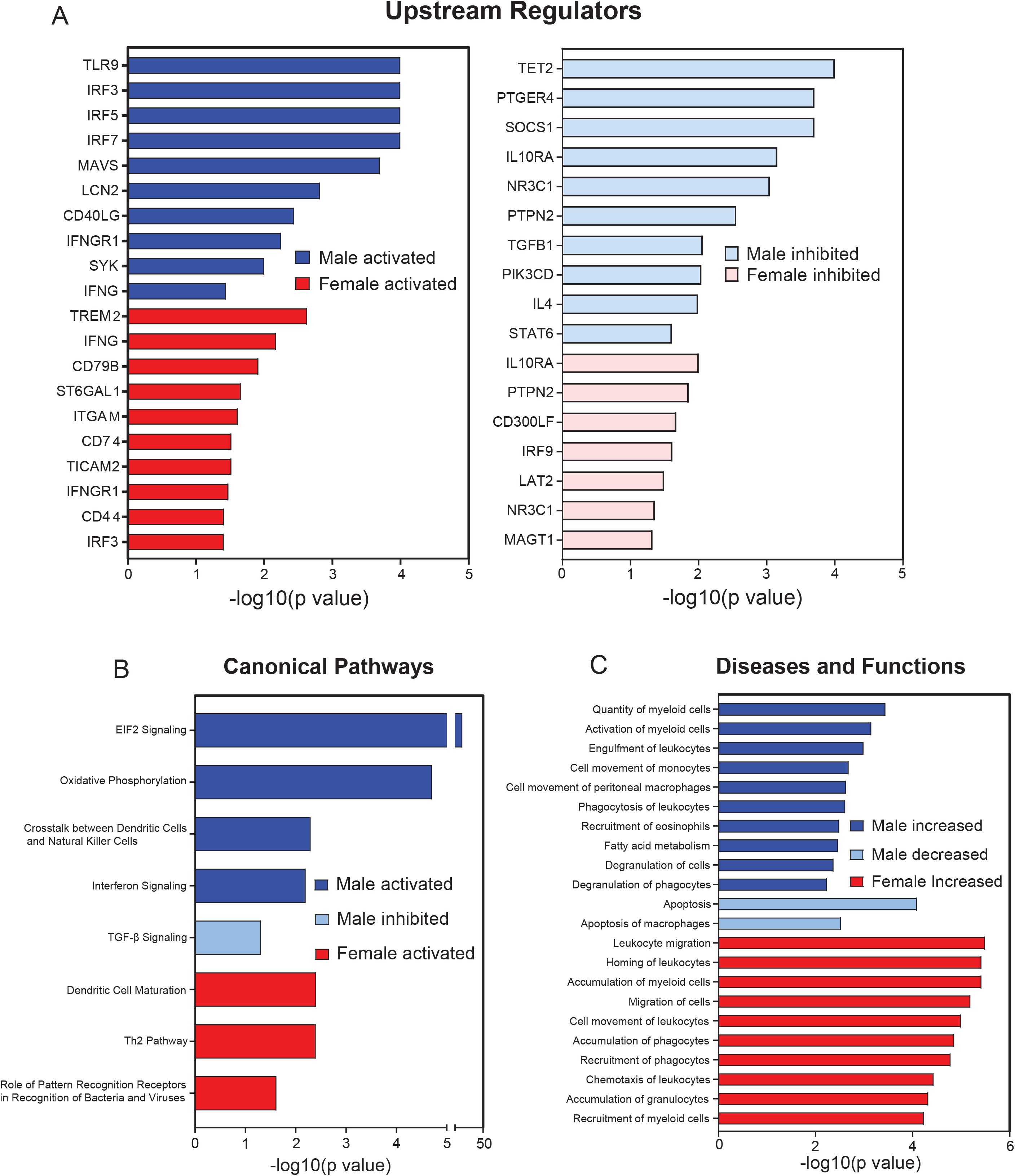
Pathway analysis reveals further sex differences in spinal microglia after CCI. (A) Upstream analysis identified top activated and inhibited candidates to regulate male and female spinal microglial gene expression after CCI. (B) The top canonical pathways activated and inhibited in male and female spinal microglia after CCI. (C) The top disease and functions altered in male and female spinal microglia after CCI. Data are shown as Benjamini-Hochberg adjusted p value.

### Microglia in the ipsilateral lumbar spinal cord have a unique, sexually dimorphic transcriptome after CCI

To better understand how the spinal microglia transcriptome after CCI compares to the transcriptome after other nerve injury models, we compared the DEGs observed in our model to two previously published nerve injury transcriptome studies describing spinal microglial transcriptome changes in male mice 7 days after spinal nerve transection (SNT; 189 DEGs upregulated) (Jeong et al., 2016) and sciatic nerve ligation (SNL; 17 DEGs upregulated) (Denk et al., 2016). When comparing the upregulated male CCI spinal microglia DEGs, there was surprisingly little overlap with the DEGs identified in the SNL and SNT lumbar spinal microglia. Specifically, only 2 genes (*Cst7* and *Lyz2*) were shared between the three groups, 8 genes (*Tspo, Olfml3, Gp2, Tmem176a, Ifi2712a* and *Ifitm3*) were shared with the SNT DEGs and 9 genes (*Cfb, Ifi30, Hcar2, Ctsl, C4b, Ccl12* and *Fcgr2b*) were shared with the SNL DEGs (**Figure 5A**). Despite the clear differences observed between microglial DEGs after different peripheral nerve injury models, we did identify some well described neuropathic pain related genes (Denk et al., 2016; Inoue and Tsuda, 2018; Jeong et al., 2016) that were upregulated in spinal microglia 7 days after CCI (**Figure 5B**). These upregulated genes appeared to be more pronounced in male microglia relative to female microglia after CCI (**Figure 5B**).

**Figure 5.**
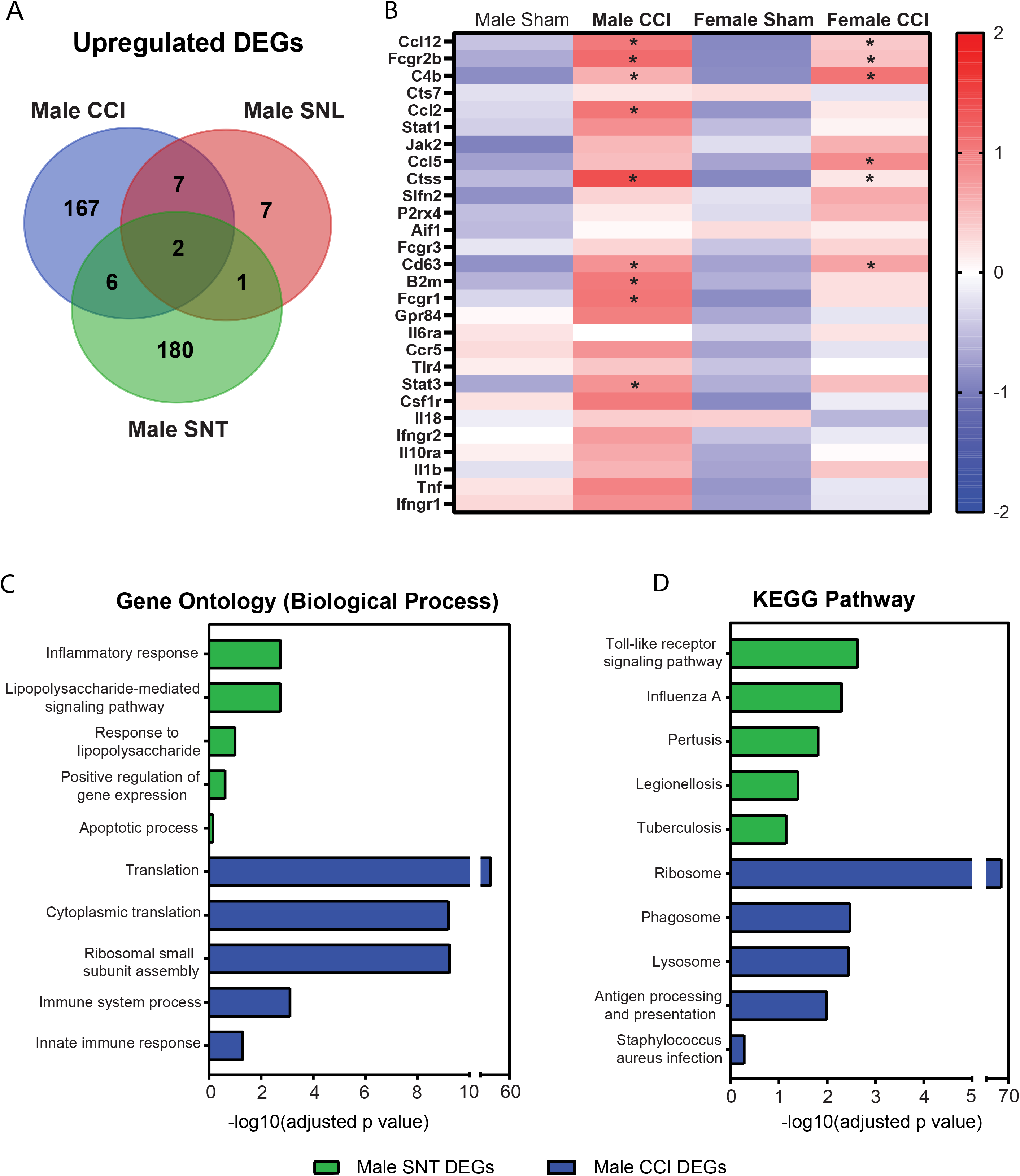
Transcriptomic differences in spinal microglia in various models of peripheral nerve injury. (A) Venn diagrams highlighting common and exclusive upregulated and downregulated DEGs found in male CCI (blue), male SNL (red) and male SNT (green) microglia in the ipsilateral lumbar spinal cord 7 days after injury. (B) Heatmap highlighting gene changes in male and female mice after CCI. Genes included are all previously described neuropathic pain related genes expressed in spinal microglia. Data are shown as z score (* p < 0.05 for CCI compared to sham). The top (C) GO enriched biological functions and (D) KEGG pathways from DEGs that are upregulated in lumbar spinal cord microglia 7 days after injury in male SNT mice (green) and male CCI mice (blue). Data are shown as Benjamini-Hochberg adjusted p value.

Gene set enrichment (GO) analysis and identification of key spinal microglia genes being discriminative between male CCI, SNL, and SNT revealed SNT DEGs are significantly involved in *inflammatory response* and *lipopolysaccharide-mediated signaling pathway* and CCI DEGs are significantly involved in *translation, cytoplasmic translation, ribosomal small unit assembly, immune system process* and *innate immune system* (**Figure 5C**). Regarding KEGG pathway enrichment, SNT DEGs are involved in *toll-like receptor signaling pathway* and *Influenza A*, whilst male CCI microglia are involved in *ribosome, phagosome, lysosome* and *antigen processing and presentation* (**Figure 5D**).

To better understand how spinal microglia transcriptome changes in response to nerve injury in comparison to the recently described DAM/MGnD and lipopolysaccharide (LPS)-stimulated microglia, we downloaded DEG list from two previously published microglia transcriptome studies (MGnD 1660 DEGs and LPS 2405 DEGs) (Keren-Shaul et al., 2017; Sousa et al., 2018) and compared them to the DEGs from male and female CCI spinal microglia (**Supplementary Table 8**). The transcription patterns from the different models demonstrated some overlap, however each signature contained unique DEGs that clearly distinguished their transcriptome. Specifically, when comparing the data with the male CCI spinal microglia DEGs, 52 upregulated genes (28%) and 6 downregulated genes (21%) were shared between the three groups, 85 upregulated genes (46%) and 7 downregulated genes (25%) were shared with the MGnD DEGs and 86 upregulated genes (47%) and 11 downregulated genes (43%) were shared with the LPS DEGs (**Figure 6A**). For the female CCI spinal microglia DEGs, 21 upregulated genes (27%) and 3 downregulated genes (18%) were shared between the three groups, 34 upregulated genes (43%) and 4 downregulated genes (24%) were shared with the MGnD DEGs and 36 upregulated genes (46%) and 7 downregulated genes (47%) were shared with the LPS DEGs (**Figure 6B**). A clear difference between these transcriptomes is that genes that are critical to microglial homeostatic function (including *Tmem119, P2ry12, P2ry13, Mef2c, SiglecH, Gpr34*) are downregulated in both LPS and DAM/MGnD models (Krasemann et al., 2017; Sousa et al., 2018) and are unchanged after CCI (**Figure 6C)**. Despite these unique differences between the microglial transcriptomes after CCI, MGnD and LPS, we did identify some critical MGnD genes that were also upregulated in spinal microglia 7 days after CCI (**Figure 6D**), including *Apoe, Axl, Grn* and *Lyz2* in both sexes as well as *Bhlhe40, Cst7, Ctsb* and *Lgals3*, exclusively in males. Again, overall changes in gene expression appeared much more pronounced in males relative to females after CCI (**Figure 6C,D**).

**Figure 6.**
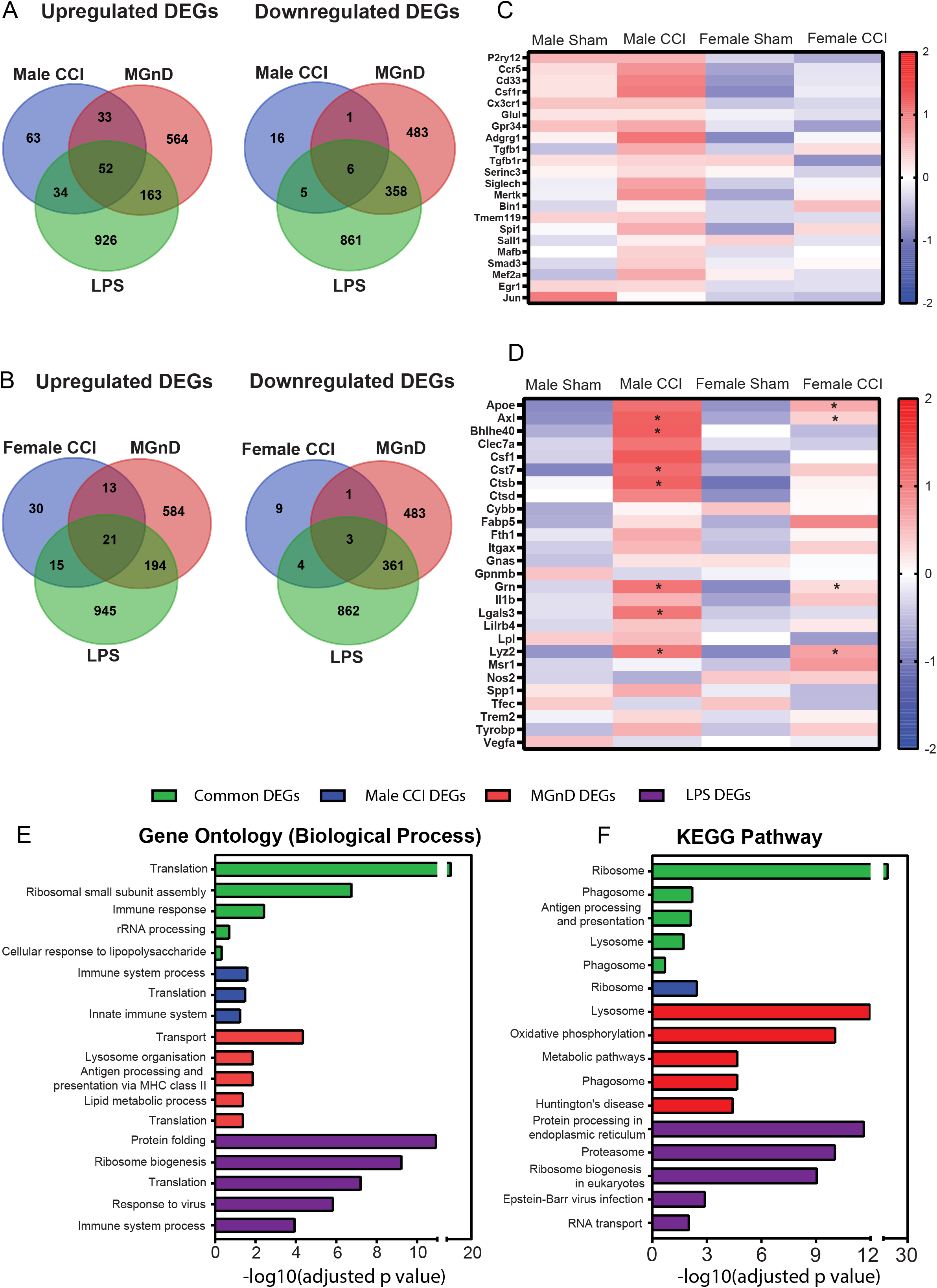
Transcriptomic differences in microglia from different disease models. (A) Venn diagrams highlighting common and exclusive upregulated and downregulated DEGs found in ipsilateral lumbar spinal cord in male mice after CCI (blue), recently described disease-associated/neurodegenerative microglia (MGnD) (red) and LPS-stimulated microglia (green). (B) Venn diagrams highlighting common and exclusive upregulated and downregulated DEGs found in female CCI (blue), MGnD (red) and LPS (green). (C and D) Heatmaps highlighting gene changes in male and female microglia after CCI. (C) Genes included are all previously described homeostatic related genes expressed in microglia. (D) Genes included are all previously described MGnD related genes expressed in microglia. Data are shown as z score (* p < 0.05 for CCI compared to sham). The top (E) GO enriched biological functions and (F) KEGG pathways from DEGs that are upregulated in male CCI, MGnD and LPS microglia (green), only in male CCI mice (blue), only in MGnD microglia (red) and only in LPS-stimulated microglia (purple). Data are shown as Benjamini-Hochberg adjusted p value.

Enrichment analysis of the shared DEGs between male CCI spinal microglia, LPS and MGnD revealed significant involvement of *translation, ribosomal small subunit assembly* and *immune response* (**Figure 6E**), with KEGG pathway activation of *Ribosome, Lysosome, Antigen processing and presentation* and *Phagosome* (**Figure 6F**). Gene set enrichment (GO) analysis and identification of key genes being discriminative between male CCI spinal microglia, LPS and MGnD revealed *translation* and *immune system process* (GO), and *ribosome pathway* (KEGG) distinguish male CCI spinal microglia, whilst high inflammatory reactivity (*protein folding and processing in endoplasmic reticulum* and *ribosome biogenesis*) distinguishes LPS and a *transport* and *lysosomal* gene signature distinguishes MGnD. There were no significantly enriched gene association with exclusively female CCI DEGs.

### Peripheral nerve injury alters microglia phenotype *in vitro*

Peripheral nerve injury activates microglia in the lumbar spinal cord, resulting in proliferation, increased volume and decreased process length and complexity (Gu et al., 2016a; Gu et al., 2016b). We examined the morphology, motility and proliferation of spinal microglia from male and female mice after CCI and sham surgery that had been polarised into homeostatic (10ng/mL macrophage colony-stimulating factor; M-CSF and 50ng/mL TGF-β) and inflammatory (10ng/mL granulocyte-macrophage colony-stimulating factor; GM-CSF) states by assessing their mean perimeter, area, sphericity, velocity and dry mass index (amount of cellular material in the field of view) over 5 hours of live cell imaging using the Livecyte microscope (**Figure 7C**). When comparing homeostatic and inflammatory spinal microglia cultures, microglia grown in GM-CSF-enriched media from both male and female spinal cord had an increase in their area and perimeter and a decrease in sphericity compared to microglia cultured in M-CSF and TGF-β-enriched media, indicative of a hypertrophied morphology (**Figure 7A** and **B**). After CCI, microglia from male mice also had an increase in their area compared to sham microglia in GM-CSF-enriched media culture (**Figure 7A**), whilst female microglia had divergent changes in sphericity after CCI in M-CSF and TGF-β-enriched media (decreased sphericity) and GM-CSF-enriched media (increased sphericity) culture, compared to sham microglia (**Figure 7B**). There was also an increase in dry mass index in microglia isolated from male CCI spinal cord compared to sham in M-CSF and TGF-β-enriched media culture, indicative of either greater proliferation or decreased apoptosis *in vitro*. There were no changes in dry mass index in female microglia cultures, or changes in motility in male and female microglia in either conditions (**Figure 7A** and **B**).

**Figure 7.**
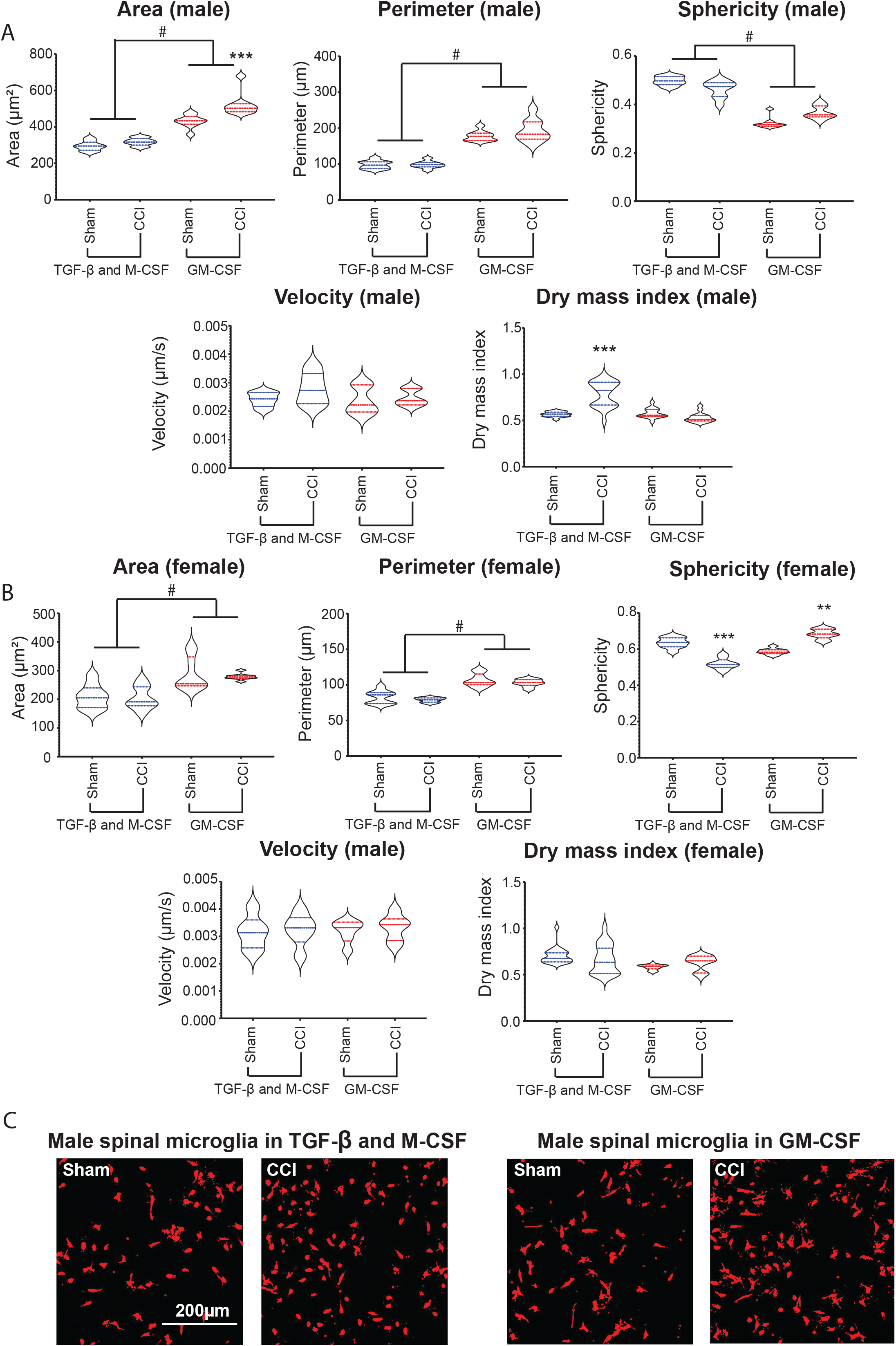
Live cell imaging of microglial phenotypes in CCI versus sham mice. The Livecyte system was used to measure the cell area, perimeter, sphericity, velocity, and dry mass index of (A) male and (B) female microglia grown in homeostatic (TGF-β and M-CSF) and inflammatory (GM-CSF) conditions. Data are shown as violin plots with median ± interquartile range (# p < 0.05 for TGF-β and M-CSF compared to GM-CSF, ** p < 0.01 and *** p < 0.001 for CCI compared to sham). (C) Representative 500um x 500um images taken from the Livecyte microscope depicting spinal microglia from male sham and CCI mice grown in homeostatic and inflammatory conditions.

### Phagocytic activity of spinal microglia *in vitro* is increased only in males after CCI

Microglial phagocytosis is crucial to the development and maintenance of neural networks (Filipello et al., 2018), and given the enrichment of lysosome and phagosome genes after nerve injury, spinal microglia may contribute to active phagocytosis. To examine phagocytic activity in male and female microglia after nerve injury *in vitro*, we measured the percentage of microglia that phagocytosed foetal calf serum (FCS)-opsonised fluorescent latex beads, which had been added to the culture media after imaging. A differential response was observed between microglia cultured under homeostatic (media enriched with M-CSF and TGF-β) and inflammatory (media enriched with GM-CSF) culture conditions (**Figure 8C**). Microglia in GM-CSF-enriched media culture had a greater phagocytic activity compared to microglia grown in M-CSF and TGF-β-enriched media culture in male mice (**Figure 8A**). Furthermore, spinal microglia obtained from CCI male mice and grown in GM-CSF-enriched media culture had a greater phagocytic capacity compared to sham spinal microglia (**Figure 8A**). In females, there were no changes in phagocytic activity in spinal microglia grown in either homeostatic or inflammatory conditions after CCI.

**Figure 8.**
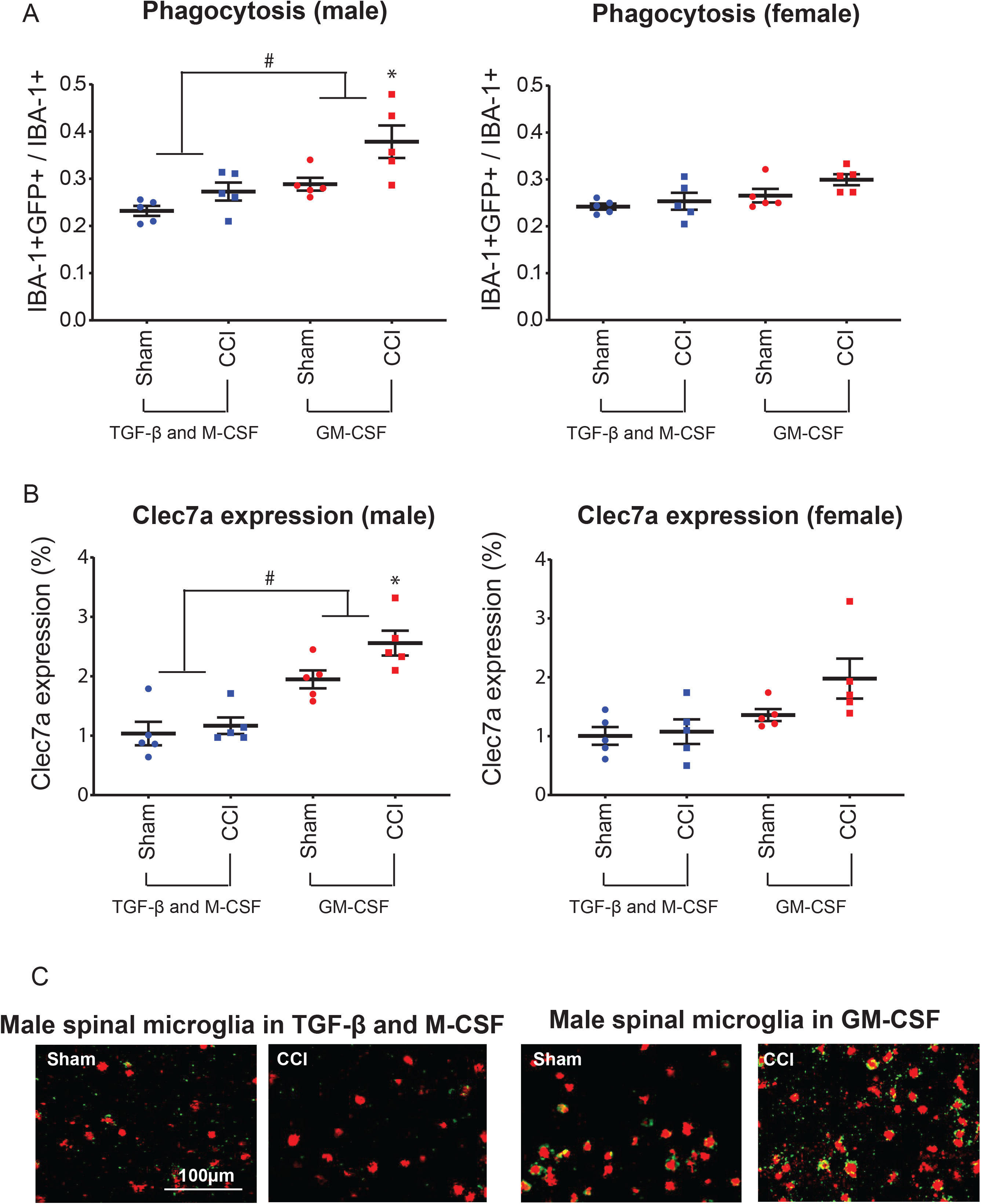
Microglial phagocytotic activity *in vitro* is greater in male mice than in female mice after CCI. (A) Latex beads were added to the microglial cultures to observe the proportion of microglia that phagocytosed the fluorescent beads in male and female microglia from sham and CCI mice grown in homeostatic (TGF-β and M-CSF) and inflammatory (GM-CSF) conditions. Data are shown as mean ± SEM (# p < 0.05 for TGF-β and M-CSF compared to GM-CSF, * p < 0.05 for CCI compared to sham). (B) Clec7a expression was measured by immunohistochemistry and compared between male and females CCI versus sham controls. (C) Representative images taken of microglia (red, IBA-1+ cells) that have engulfed fluorescent latex beads (green) from male sham and CCI mice, grown in homeostatic and inflammatory conditions.

We next measured the expression of C-lectin domain containing 7a (Clec7a), a transmembrane protein involved in phagocytosis, which was significantly upregulated in microglia cultured in GM-CSF compared to microglia grown in M-CSF and TGF-β cultured media (**Figure 8B**). After CCI, male spinal microglia had a higher expression of Clec7a compared to sham microglia when cultured in GM-CSF-enriched media (**Figure 8B**). Again, there were no changes in Clec7a expression in female spinal microglia grown in either of the culture conditions after CCI.

## DISCUSSION

Neuroimmune sex-dependent differences in pain processing are well recognized, however the contribution of microglia to neuropathic pain in males and females remains controversial. Here, we investigated the microglial transcriptome in spinal and supraspinal pain related regions in different models of peripheral neuropathic pain (CCI of the sciatic nerve and paclitaxel- and oxaliplatin-induced peripheral neuropathy). We show that there is no common microglia signature across the different CNS regions and the different models of neuropathic pain. In addition, a substantial transcriptomic change was only observed in microglia within the ipsilateral lumbar spinal cord after peripheral nerve injury. Specifically, both male and female mice upregulate genes associated with inflammation, phagosome, and lysosome activation 7 days after CCI compared to sham control mice. Interestingly, microglia isolated from male mice after CCI revealed a prominent global transcriptional shift compared to female mice, reflecting highly activated cells. In addition, *in vitro* studies revealed that only spinal microglia from male mice are more prone to develop a reactive phenotype and phagocytose debris after CCI in inflammatory culture conditions.

Our results show that despite all neuropathic pain models inducing mechanical allodynia and exploratory behavioural changes, no common microglial signature in the spinal cord or supraspinal regions could be identified. Moreover, we found only few changes in microglial gene expression in pain-related regions in our CIPN models and in supraspinal regions after CCI. There is an ongoing debate on the involvement of microglia in CIPN, with some studies identifying astrocytes rather than microglia as critical in the development of neuropathic pain in CIPN (Makker et al., 2017; Robinson et al., 2014; Zhang et al., 2012; Zheng et al., 2011). Interestingly, previous studies have identified acute activation (7 days after first injection) of spinal microglia in response to paclitaxel and oxaliplatin treatment that dissipates over time (Mannelli et al., 2014; Ochi-ishi et al., 2014) and microglial activation in a methotrexate model of CIPN is critical in shifting astrocytes to a reactive phenotype (Gibson et al., 2019). Taken together, while we cannot dismiss a role for microglia in the early development of CIPN, our data highlights that there are no ongoing gene changes in microglia in the paclitaxel and oxaliplatin models of CIPN.

Microglial activation in brain regions is important for mediating the affective-motivational and cognitive dimensions of neuropathic pain (Austin and Fiore, 2019; Bushnell et al., 2013). Despite this, it is unknown whether the mechanism of microglia activation is region dependent or common throughout the CNS. Here, we observed a substantial transcriptomic change in microglia within the ipsilateral lumbar spinal cord and minor gene transcript changes in supraspinal pain-related regions after CCI. Microglial changes in supraspinal regions have previously been observed in rodents within 14 days following CCI (Fiore and Austin, 2019; Mor et al., 2010; Taylor et al., 2017), though a recent study identified that microglia are activated in specific pain-related regions in mice only at delayed time points after CCI, concurrent with the presence of affective-motivational behavioural changes (Barcelon et al., 2019). There is also evidence that microglial alterations in the hippocampus oppose those observed in the spinal dorsal horn after spared nerve injury in rats (Liu et al., 2017). This implies that supraspinal microglia activation occurs in a different timeframe to spinal microglia and development of pain behaviours and is likely region-dependent.

It remains contentious whether there are sexually dimorphic differences in the contribution of spinal microglia to pain after nerve injury (Mogil, 2020). It has been proposed that microglia are critical for nerve injury-induced hypersensitivity in male mice, particularly via the P2RX4-BDNF-TRKB pathway and activation of p38 mitogen-activated protein kinase (MAPK) (Masuda et al., 2014; Tsuda et al., 2004; Tsuda et al., 2003), whilst in female mice inhibition of these microglial pathways is ineffective in reducing pain hypersensitivity (Sorge et al., 2015; Taves et al., 2016). Further, females appear to be dependent on T cells to mediate pain hypersensitivity following nerve injury, though the detailed mechanisms of this process remain controversial (Lopes et al., 2017; Sorge et al., 2015). However, there are numerous studies that report no obvious sex differences in the suppressive effect of microglial inhibitors, genetic knockout of microglial-selective molecules or ablation of microglia in various nerve injury models (Barragan-Iglesias et al., 2014; Gu et al., 2016a; Peng et al., 2016; Staniland et al., 2010). Our results show that spinal microglia from both male and female mice 7 days after sciatic nerve CCI are characterised by an increase in upregulated genes involved in microglial activation and inflammatory immune response (including *Ifitm3, Axl, H2-q7, Ly86, Jak3, H2-d1, C4b, Tspo, Gas6, Ccl12*), as well as lysosome (including *Cd63, Gm2a, Lamp1, Ctsl, Ctsh, Man2b1, Ctss, Ctsb*) and phagosome (including *Itgam, Lamp1, Fcgr4, Ctsl, H2-q7, Fcgr2b, Ctss, H2-d1*) activation. It is particularly surprising that no changes in *Csf1r, Tyrobp, P2rx4, Irf5* or *Irf8* expression were identified in either sex, as upregulation of these genes has been previously reported as critical for the development of mechanical hypersensitivity following nerve injury (Guan et al., 2016; Masuda et al., 2014; Masuda et al., 2012). Previous studies identified transcriptomic activation of microglia and enrichment of interferon-γ (IFNγ) and fragment-crystallizable γ receptors (FcγRs) signalling, as well as lysosome activation (Denk et al., 2016; Franke et al., 2016; Jeong et al., 2016; Tsuda et al., 2009). In our data, pathway analysis identified the interferon-related transcription factors IRF3, IRF7 and IRF5 as upstream mediators of CCI-induced gene regulation in males and IFNγ along with its receptor IFNGR1 in both male and female microglia as important upstream mediators.

Regarding sex-specific changes, we found that the majority of DEGs in lumbar spinal cord are sex specific (Male 79% and Female 53%). Interestingly, microglia isolated from male mice after CCI revealed a prominent global transcriptional shift compared to female mice, displaying an over representation of ribosome/translation genes as well as microglial activation and inflammatory immune response (including *Fcgr1, Irf7, B2m* and *Bhlhe40*), thus reflecting a highly activated transcriptome compared to female microglia. This transcriptional shift was supported by pathway analysis, which identified the chronic pain-related translation regulation signalling pathway EIF2 (Khoutorsky et al., 2016) as the most active canonical pathway in males (**Figure 4**). On the other hand, CCI microglia from female mice had an increase in genes involved in transport (including *Apoe* and *Grin2b*). Pathway analysis identified *Trem2* as the top upstream mediator of CCI-induced gene regulation in female microglia (**Figure 4A**). Activation of the 12-kDa transmembrane protein (DAP12)-dependent signalling by TREM2 is critical to the development of pain after nerve injury (Guan et al., 2016) and apolipoprotein E (APOE) activation of TREM2 is critical to the development of MGnD (Krasemann et al., 2017), whereas *Trem2* gene is downregulated in microglia after LPS treatment (Sousa et al., 2018).

In comparison with other studies that investigated spinal microglial transcriptomes after nerve injury, there were only 2 common genes (*Cst7* and *Lyz2*) upregulated in male microglia after CCI, SNT and SNL. *Cst7* and its encoding protein Cystatin F are markers of ongoing demyelination with concurrent myelination (Ma et al., 2011), a process which is important for the development of neuropathic pain (Chu et al., 2020) and *Lyz2* (Lysozyme C-2) is also found in regions of acute demyelination, where it plays a role in the intracellular sorting of major histocompatibility complex class II (Plemel et al., 2020). Surprisingly, there was more gene signature overlap with the MGnD phenotype (e.g., *Apoe, Axl, Bhlhe40, Lgals3, Cst7, Ctsb, Grn, Lyz2*) (**Figure 6D**). Notably, previous studies have revealed that *Lgals3* (Galectin 3) inhibition attenuates pain after nerve injury (Ma et al., 2016), *Grn* (Granulin) promotes peripheral nerve regeneration (Lim et al., 2012), *Apoe* polymorphisms are related to severity of peripheral neuropathy (Monastiriotis et al., 2013; Monastiriotis et al., 2012) and *Ctsb* (Cathepsin B) inhibition attenuates development of allodynia in inflammatory pain (Ma et al., 2016; Sun et al., 2012). *Axl* (AXL Receptor Tyrosine Kinase) signalling via its ligand growth arrest gene 6 (*Gas6*) that is also upregulated after CCI, facilitates phagocytosis of apoptotic cells by microglia as well as increasing cellular migration and myelination and reducing apoptosis and toll-like receptor-mediated inflammation (Gilchrist et al., 2020; Goudarzi et al., 2020; Grommes et al., 2008). *Bhlhe40* is a transcription factor that regulates myeloid cell activation and inflammation as well as self-renewal (Carey et al., 2020; Jarjour et al., 2019). There was substantially less overlap when comparing downregulated genes in the MGnD phenotype (25% in males, 24% in female). Importantly, there was no downregulation of homeostatic genes that are a hallmark of impaired TGF-β signalling and development of MGnD and DAM phenotypes (Krasemann et al., 2017). Contrastingly, there was a trend of increased expression of these genes in males after CCI (**Figure 6A**). These data highlight that not only are the gene signatures expressed by male and female spinal microglia following CCI sexually distinct, but they are unique when compared to other microglial signatures from nerve injury models and CNS diseases.

To our knowledge, live cell imaging of primary microglia isolated from nerve-injured mice has not been performed previously. Our *in vitro* studies revealed that only spinal microglia from male mice are more prone to develop a reactive phenotype and phagocytose debris after CCI in inflammatory (GM-CSF) culture conditions. An increase in microglial area in culture is reflective of a shift towards a more reactive phenotype (Caldeira et al., 2014), indicating that CCI may prime microglia to become more reactive to an inflammatory insult (Kinuthia et al., 2020). The increase in dry mass index is unlikely to be due to proliferation *in vitro*, rather this may reflect a reduction in cell death potentially due to the inhibition of apoptosis in male microglia after CCI (**Figure 4C**).

Microglia are reported to increase their phagocytic activity in the spinal cord 7 days after nerve injury (Echeverry et al., 2008; Nishihara et al., 2020). Indeed, our RNA-seq analysis revealed an upregulation of phagosome-related genes in spinal microglia after CCI and identified increased phagocytosis by pathway analysis (**Figure 4C**). Using phagocytosis assay *in vitro*, we observed CCI-induced increases in phagocytic activity and CLEC7A expression in male microglia grown in GM-CSF. We did observe a non-significant increase in *Clec7a* gene expression (1.75-fold increase, unadjusted p = 0.07) in our RNA-seq data which was not present in female mice (0.1-fold decrease, unadjusted p = 0.55). CLEC7A-mediated phagocytosis has been reported to be a critical feature of the MGnD phenotype, and microglia isolated after CNS injury and cultured *in vitro* have increased phagocytic activity (REF) (Fu et al., 2014; Fumagalli et al., 2019), further highlighting the overlap between MGnD and the microglial phenotype observed in male spinal microglia after CCI. Whilst microglial sex differences in primary culture haven’t been investigated in nerve injury models, sex differences have been observed in GM-CSF enriched microglial cultures in a model of ischemic stroke (Bodhankar et al., 2015). These data also support the idea of differential reactivity in microglia between the two sexes even when grown in primary cultures (Villa et al., 2018).

In summary, our findings present a comprehensive transcriptomic view of microglia involvement throughout the neuroaxis in different models of peripheral neuropathy, and reveal a lack of common neuropathic gene signature, as well as the presence of sex differences between male and female spinal microglia in neuropathic pain states. Our findings suggest that new sex-specific therapeutic approaches are needed to restore detrimental microglial phenotypes found in neuropathic pain. A potential limitation of our study is that we used bulk microglia for RNA-seq analysis, as well as pooling microglia from pain-related CNS regions. It is possible that differential and distinct activation status existed at individual cell level. In addition, multiple CNS cell types communicate and mutually depend on each other to function. The activity of microglia is especially linked to astrocyte function. For example, it was recently shown that microglia activation induces neurotoxic reactive astrocyte formation that contributes to cognitive impairment in a CIPN model (Gibson et al., 2019). Therefore, a comprehensive study of the molecular changes in different single-cell types, together with bioinformatics tools, are needed to further our understanding of sexual dimorphism in neuropathic pain and identify novel sex-specific therapeutic targets.

## Methods

### Animals

All experiments were conducted on adult (8-12 weeks old) male and female C57BL/6 mice (Australian BioResources, Moss Vale, NSW, Australia) with a value of 6 per group unless otherwise stated. Animals were housed in ventilated standard cages with free access to food and water and maintained on a 12:12-h light/dark cycle. Mice were acclimatised to the animal facility for at least one week prior to experiments commencement. All behavioural tests were conducted during the dark cycle. All animal experiments were approved by the Animal Care and Ethics Committee of the University of New South Wales (UNSW) Sydney, Australia.

### Peripheral nerve injury model

Chronic constriction injury (CCI) was performed on the left sciatic nerve under anaesthesia (4% iosflurane in oxygen for the duration of surgery) by loosely tying two chromic gut ligatures (6-0 Ethicon) around the sciatic nerve proximal to the trifurcation 1 mm apart to impair but not arrest epineural blood flow. Corresponding sham surgery exposed the sciatic nerve but no ligatures were tied. The muscle layers were closed with sutures (Mersilk 5-0 Ethicon) and skin closed with Michel clips (9mm BD Diagnostics). Mice were monitored daily after surgery.

### Chemotherapy-induced peripheral neuropathy model

CIPN mouse models were carried out with two different chemotherapeutic drugs: paclitaxel and oxaliplatin. Paclitaxel (In vitro Technologies) was dissolved in 600 µl absolute ethanol per 10mg to make a stock solution of 16.66 µg/µl. The stock solution was further diluted using a 1:1:8 ratio of paclitaxel, cremaphor and 0.9% sterile saline, respectively. The vehicle control solution was prepared similarly, excluding paclitaxel. Oxaliplatin (Sigma-Aldrich) was dissolved in sterile 5% dextrose/water to a stock solution of 1mg/mL and stored at -30°C until the day of treatment. In each experiment, the mice were randomly assigned into 4 groups: vehicle paclitaxel control, paclitaxel, vehicle oxaliplatin control and oxaliplatin. Each mouse was weighed prior to each injection and injected intraperitoneally (i.p) with the appropriate volume for their treatment, using the standard of 5 mg/kg for paclitaxel (or vehicle paclitaxel control) and 2.5 mg/kg for oxaliplatin (or vehicle oxaliplatin control). A total of 6 courses of injections were given at days 0, 2, 4, 6, 8 and 10, resulting in a cumulative dose of 30 mg/kg for paclitaxel and a total of 12 injections were given at day 0, 1, 2, 3, 7, 8, 9, 10, 14, 15, 16, 17 resulting in a cumulative dose of 30 mg/kg for oxaliplatin.

### Behavioural tests

#### Von Frey testing

Mechanical sensitivity was assessed prior to nerve injury or initial paclitaxel, oxaliplatin or vehicle control injection and 5-days after nerve injury or last injection during the dark phase by the up-down method using calibrated von Frey filaments. Mice were habituated to the behavioural testing apparatus for at least 30minutes before data collection in a quiet and well-controlled environment. Mechanical withdrawal threshold was assessed using the up-down method. Briefly, mice were placed into the test cage with an elevated mesh, and stimulating the mid-plantar surface using a set of 8 calibrated von Frey filaments (0.02, 0.04, 0.07, 0.16, 0.4, 0.6, 1.0 and 1.4g) until the filament bent slightly. A positive response was recorded when a withdrawal reflex was observed. The first filament was always 0.4g and if there was a positive response, the 0.16g filament was applied and if there was a negative response the 0.6g filament was applied. Each hindpaw was tested 4 times after the initial positive response, and the 50% paw withdrawal threshold for each hindpaw was calculated. The interval between trials on the same paw was at least 3 minutes. For the CIPN models, the left- and right-hand paws were averaged.

#### Gait assessment

For mice undergoing paclitaxel, oxaliplatin and relevant vehicle controls, motor coordination was examined using the DigiGait system (Mouse Specifics Inc, Quincy, MA, USA). Baseline was established prior to initial injection and testing was performed 5-days after the last injection. Briefly, the testing apparatus consisted of a motorised treadmill with a transparent belt and imaged from beneath with a high-speed digital video camera to capture the paw prints on the belt. Prior to testing, mice were first habituated to the treadmill for 5mins. Before recording, the mice were given 30s to run at 15 cm/s after which a 5s recording of continuous running was taken. For each 5s recording, the paws were identified, and background was reduced to remove the snout and tail from the analysis. Images were collected at a rate of 140 frames/s and stored as audio video interleaved (AVI) files for later analysis. The analysis was automated by the mouse specific analysis software as part of the DigiGait system package. The paw values were averaged per mouse and a cumulative gait index was calculated as previously described (Lambert et al., 2014).

#### Open field activity test

Exploratory behaviour testing was performed 6-days after nerve injury or last chemotherapy injection using a photobeam activity system (PAS; San Diego Instruments, San Diego, CA) in a climate-controlled room. Mice were placed in the centre of a 40 cm (width) × 40 cm (diameter) × 38 cm (height) open-top PAS chamber surrounded by a customized open-top box made of white Perspex occluding vision of the surrounding room except for the ceiling. Nose-poke and rearing events were recorded for 5 minutes by quantifying beams breaks. To measure nose-poke, we used a manufacturer supplied flooring containing 16 evenly spaced nose-poke holes (hole-board), which were laser activated each time the mice investigated the holes. Beam break recordings were processed using the manufacturer’s software to give quantitative distance travelled, average speed, time spent in the centre of the field, nose-poke and rearing data.

### Flow cytometry microglial sorting

For microglial cell sorting, mice were deeply anaesthetised with sodium pentobarbital (200mg/kg i.p.) and then transcardially perfused with 50mL hanks balanced salt solution (HBSS) containing heparin (1:1000). Following perfusion, four blocks of CNS were dissected out following CCI or Sham surgery using the Allen mouse brain atlas; (1) the ipsilateral (left) lumbar spinal cord (L3-L5); (2) the medial prefrontal cortex, hippocampus and amygdala; (3) posterior thalamus and S1 cortex relating to the hindlimb; and (4) the periaqueductal gray and rostroventral medulla. For the CIPN treatments, two blocks were dissected out containing the lumbar spinal cord (L3-L5), and the medial prefrontal cortex, hippocampus and amygdala block. Each block of tissue was homogenised using a dounce glass tissue homogeniser. Mononuclear cells were separated through Percoll (GE Healthcare Life Sciences) 37%/70% gradient centrifugation. Mononuclear cells were isolated from the interface and stained on ice for 30 min with combinations of BV650 rat anti-mouse CD11b (marking myeloid cells; 1:300), FITC rat anti-mouse Ly6C (marking monocytes/macrophages; 1:300) and APC rat anti-mouse 4D4 (marking resident microglia; 1:1000) in blocking buffer containing 0.2% bovine serum albumin (BSA, Sigma-Aldrich) in HBSS. Cell sorting was performed using FACSAriaIII cell sorter (Becton Dickson). Microglial cells were identified as CD11b+ Ly6C-4D4+ and sorted directly in 1.5mL Eppendorf tubes and stored at -80°C.

### Bulk RNA-seq

Bulk RNA sequencing was performed as previously described (Butovsky et al., 2014). Briefly, 1,000 isolated Ly6C^-^CD11b^+^4D4^+^ microglia were lysed in 5ul TCL buffer + 1% β-mercaptoethanol. Smart-Seq2 libraries were prepared and sequenced by the Broad Genomic Platform. cDNA libraries were generated from sorted cells using the Smart-seq2 protocol^5^. RNA sequencing was performed using Illumina NextSeq500 using a High Output v2 kit to generate 2 × 38 bp reads. Transcripts were quantified using Salmon v1.4. Raw read counts were processed and normalized in R using DESeq2’s median of ratio method and low abundance genes were filtered below a mean count of 5 reads/sample. DEGs were called using the DESeq2 program in R (v3.6.3) with a Benjamini-Hochberg adjusted p-value < 0.05.

### RNA-seq data processing and network analysis

Network analysis was conducted using Database for Annotation, Visualisation and Integrated Discovery (DAVID) and Ingenuity pathways analysis (IPA, Qiagen). Briefly, DEGs (with corresponding fold changes and p values) were incorporated in canonical pathways and bio-functions were used to generate biological networks. DAVID was performed to identify gene ontology (GO) enrichment analysis to identify GO biological process terms that are over-represented and KEGG pathway analysis to identify molecular pathways and biological functions that are likely to be encoded in the genome. IPA was performed to identify most significant canonical pathways, diseases and functions, and gene networks and to categorize DEGs in specific diseases and functions. Gene networks and pathways with an adjusted p < 0.05 and a z score > 2 or < -2 were considered significantly upregulated or downregulated, respectively. Custom Venn diagrams were calculated and drawn with an online tool (Bioinformatics and Evolutionary Genomics: bioinformatics.psb.ugent.be/webtools/Venn/).

### Real-time PCR

For qRT-PCR validation of our above RNA-seq data, we FACS-sorted the microglia as described above, using only ipsilateral lumbar spinal microglia after sham and CCI in male and female mice. Briefly, microglia samples were pooled from two mice and then lysed in RLT buffer with beta-mercaptoethanol, and RNA was extracted using QIAGEN RNeasy Micro Kit following manufacturers instructions. RNA quantity and quality was assessed on DeNovix DS-11 Spectrophotometer. Total RNA (40ng) was used in 20 µL of reverse transcription reaction (SuperScript IV VILO Master Mix with ezDNase Enzyme kit, Invitrogen) according to the manufacturer’s instruction and 2 µL of cDNA in 20 µL reverse transcription reaction with specific FAM-labelled Taqman probes (*Apoe* Mm01307193_g1, *Axl* Mm00437221, *B2m* Mm00437762_m1, *Bhlhe40* Mm00478593_m1, *Ccl12* Mm01617100_m1, *Csf1r* Mm01266652_m1, *Cst7* Mm00438351, *Fcgr2b* Mm00438875, *P2ry12* Mm01950543_s1, *Gapdh* Mm9999915_g1, *Rps18* Mm02601777_m1, Cat#4453320 Thermofisher). We used the ddCT method, normalising each sample to the average of GAPDH and the same sex sham microglia pool. All qRT-PCRs were performed in duplicate presented as mean ddCT ± SEM.

### Primary adult mouse microglial culture

Sorted microglia from CCI or sham injured spinal cord were cultured in 96-well plate (1.5 × 10^4^ cells per well in 0.2mL) in poly-D-lysine-coated (Sigma-Aldrich) plates and grown in microglia culture medium (DMEM/F-12 Glutamax, ThermoFisher) supplemented with 10% foetal calf serum (FCS, Sigma-Aldrich), 100 U/mL penicillin (Sigma-Aldrich, 100 U/mL streptomycin (Sigma-Aldrich) at 37 °C and 5% CO_2_. Microglia were polarised to reflect CNS homeostatic or inflammatory conditions with additional cytokines. To generate ‘homeostatic’ microglia, sorted microglia were cultured in microglia culture medium containing recombinant carrier-free M-CSF 10ng/mL and 50ng/mL human recombinant TGF-β for 5 days. To generate ’inflammatory’ microglia, sorted microglia were cultured in microglia culture medium containing recombinant carrier-free GM-CSF 10ng/mL. The plates were stored in the incubator at 37 °C and 5% CO_2_ for five days before imaging.

### Live Cell Imaging

After five days of primary spinal microglia cultures, the cultured media was replenished with equivalent media and cytokines. Live cell imaging was then performed using the Livecyte microscope system (Phasefocus). Microglia were then imaged at 20x objective every 20 minutes and tracked for 5 hours on the Livecyte microscope system. Each well had duplicate 500µm x 500µm regions and the Livecyte system segmentation analysis (Phasefocus) was used to track changes in proliferation (dry mass index), morphology (cell area, perimeter and sphericity) and motility (mean velocity) of the microglia in culture.

### Phagocytosis Assay and Immunocytochemistry

Aqueous green fluorescent latex beads of 1µm diameter (Sigma-Aldrich, L1030) were pre-opsonised in FCS in a ratio of 1:5 for 1 hour at 37°C. The beads containing FCS were diluted with microglia culture medium without 10% FCS to reach a final concentration of 0.01% beads and 0.05% FCS. The media in the wells were removed and replaced with 100µL of phagocytosis assay media. The plate was incubated at 37°C for 1 hour, and then the cultures were washed thoroughly with PBS and fixed with 4% paraformaldehyde for 15 minutes. The fixed microglia were permeabilised with 0.1% Triton X-100 in PBS for 15 minutes and blocked with 5% Normal Donkey Serum (NDS; Sigma-Aldrich, D9663) in 0.1% Tween in PBS (PBS-T) for 30 minutes. The microglia were incubated with rabbit anti-mouse IBA-1 (ionised calcium-binding adapter molecule 1; macrophages/microglia; 1:500; Wako Chemicals) and rat anti-mouse Clec7a (activated microglia; 1:50; InvivoGen, CA, USA) in 2% NDS and PBS-T for 1 hour at 4°C. The microglia were washed three times with PBS. The microglia were incubated with the secondary antibodies, Alexa Fluor 488 donkey anti-rabbit (1:500 IgG; Life Technologies, A-21206), and Alexa Fluor 594 donkey anti-rat (1:500 IgG; Life Technologies, A-21209) in 2% NDS and PBS-T for 30 minutes. The microglia were washed, then incubated with Hoechst 33342 (nuclei stain; 1:10,000; Life Technologies) for 15 minutes, before being washed and stored with PBS in each well at 4°C in the dark until imaging.

### Imaging and analysis

The ZEISS LSM 900 (ZEISS Australia) was used to acquire images for analysis of Clec7a, IBA-1 expression and GFP^+^ beads. Images were acquired using 10x objective and the entire well containing primary microglia was imaged at 512×512 resolution using the tile scan function. All image analysis was then completed using ImageJ (Bethesda, USA). Each image was converted to 8-bit grey-scale and threshold adjusted. The area of red pixels was measured as a percentage of the total area of the image for statistical analysis of IBA-1 and Clec7a protein expression. The phagocytic activity of the polarised microglia was determined by calculating the percentage of microglia containing beads. The total numbers of microglia (IBA-1^+^ cells) and IBA-1^+^ cells containing beads (IBA-1^+^GFP^+^) were then manually counted using ImageJ, and expressed as a percentage of IBA-1^+^ cells.

### Statistical analysis

Statistical analyses were conducted using GraphPad Prism 9.0.0 (Von Frey, Open field, DigiGait, Live cell imaging and Phagocytosis assay), DAVID (GO and KEGG pathways) and IPA (Upstream regulator, Canonical pathways, Disease and functions). Von Frey, gait analysis and live cell imaging data were analysed using a two-way mixed effects analysis of variance (ANOVA) with sidaks multiple comparisons test. Open field and qRT-PCR analysis was performed using an unpaired t test between CCI or CIPN groups and their respective controls for each sex. Phagocytosis assay analysis and Clec7a immunofluorescence were analysed using a one-way ANOVA with Tukey’s multiple comparisons test. The criterion for significance was p < 0.05 for all analyses unless otherwise stated.

## Acknowledgements

This study was supported by a grant from the National Health and Medical Research Council of Australia awarded to GMT and OB (ID # APP1162060), and partially by the Cancer Institute NSW Translational Program Grant – “Chemotherapy-induced Peripheral Neuropathy: Assessment strategies, Treatment and Risk Factors” (ID # 14/TPG/1-05) (GMT). This work was also supported by the NIH-NINDS (1R01NS088137) (O.B.), NIH-NIA (R01AG051812, R01AG054672) (O.B.), the Cure Alzheimer’s Fund (O.B.) and BrightFocus Foundation A2021022S. The funders had no role in study design, data collection and analysis, decision to publish or preparation of the manuscript. We thank the Mark Wainwright Analytical Centre, in particular the Flow Cytometry Facility and the Biomedical Imaging Facility at UNSW Sydney, Australia and the Broad Institute of MIT and Harvard Cambridge, USA.

## Author Contributions

N.F.T., G.M-T., and O.B. designed research; N.T.F. performed experiments, analysed data; N.T.F., Z.Y., D.G., C.D.G., Z.Y., and A.D. performed RNaseq analysis, k-means clustering and network analysis; N.T.F., Z.Y., J.H., microglia culture and phagocytosis assay; N.T.F. wrote the first draft of the paper; N.T.F and G.M-T wrote the paper. All authors discussed the results and conclusions and reviewed the manuscript.

## Figure Legends

**Supplementary Figure 1. Microglia isolation with gating strategy**.

**Supplementary Figure 2. qRT-PCR validation of gene changes after CCI**.

**Supplementary Table 1. Comparison of DEGs in spinal microglia after CCI between males and females**.

**Supplementary Table 2. Top upstream regulators identified in male spinal microglia after CCI by ingenuity pathway analysis (IPA)**.

**Supplementary Table 3. Top upstream regulators identified in female spinal microglia after CCI by ingenuity pathway analysis (IPA)**.

**Supplementary Table 4. Top canonical pathways identified in male spinal microglia after CCI by ingenuity pathway analysis (IPA)**.

**Supplementary Table 5. Top canonical pathways identified in female spinal microglia after CCI by ingenuity pathway analysis (IPA)**.

**Supplementary Table 6. Top disease functions identified in male spinal microglia after CCI by ingenuity pathway analysis (IPA)**.

**Supplementary Table 7. Top disease functions identified in female spinal microglia after CCI by ingenuity pathway analysis (IPA)**.

**Supplementary Table 8. Comparison of DEGs in spinal microglia after CCI compared to Neurodegenerative and LPS stimulated microglia**.

---

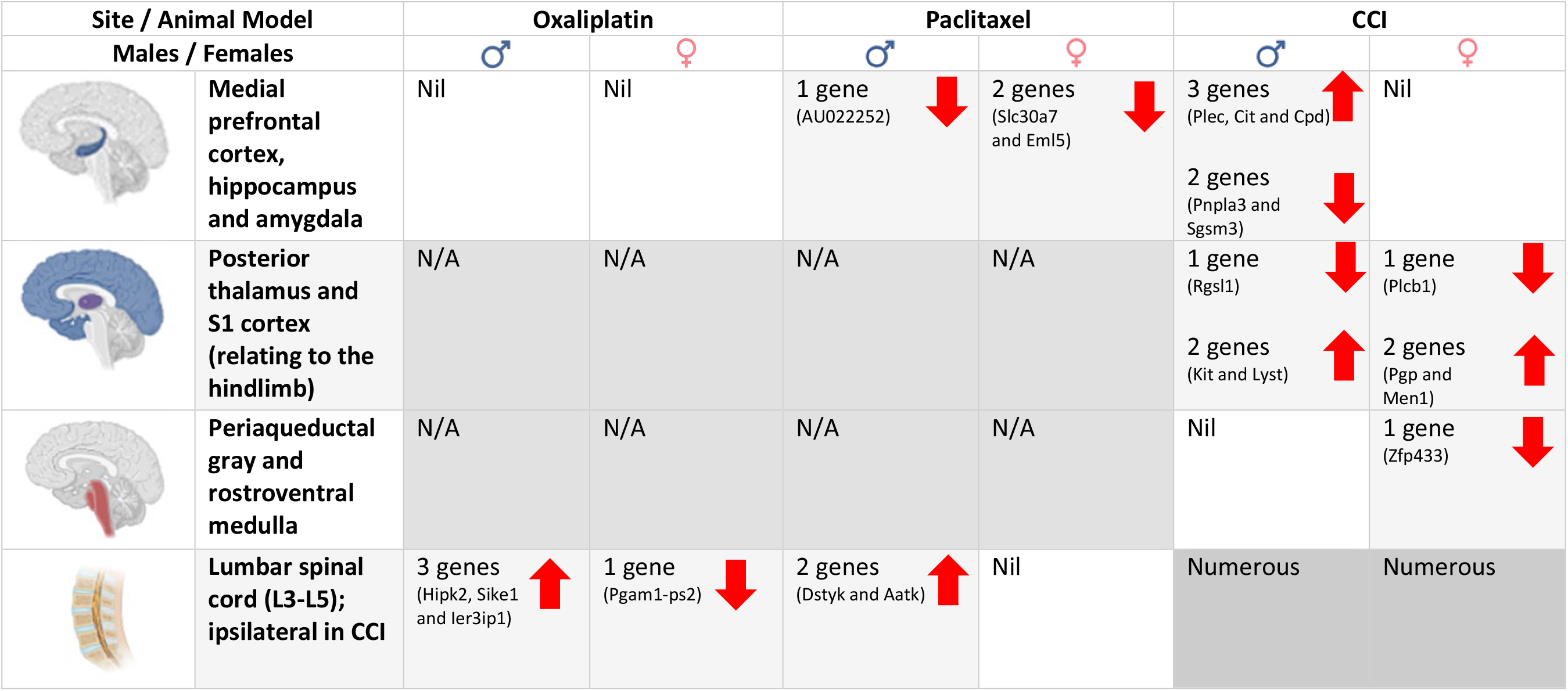

## References

Austin, P.J., and Fiore, N.T. (2019). Supraspinal neuroimmune crosstalk in chronic pain states. Current Opinion in Physiology 11, 7–15.

Austin, P.J., and Moalem-Taylor, G. (2010). The neuro-immune balance in neuropathic pain: involvement of inflammatory immune cells, immune-like glial cells and cytokines. J Neuroimmunol 229, 26–50.

Barcelon, E.E., Cho, W.H., Jun, S.B., and Lee, S.J. (2019). Brain Microglial Activation in Chronic Pain-Associated Affective Disorder. Front Neurosci 13, 213.

Barragan-Iglesias, P., Pineda-Farias, J.B., Cervantes-Duran, C., Bravo-Hernandez, M., Rocha-Gonzalez, H.I., Murbartian, J., and Granados-Soto, V. (2014). Role of spinal P2Y6 and P2Y11 receptors in neuropathic pain in rats: possible involvement of glial cells. Mol Pain 10, 29.

Blyth, F.M., March, L.M., Brnabic, A.J., and Cousins, M.J. (2004). Chronic pain and frequent use of health care. Pain 111, 51–58.

Bodhankar, S., Lapato, A., Chen, Y., Vandenbark, A.A., Saugstad, J.A., and Offner, H. (2015). Role for microglia in sex differences after ischemic stroke: importance of M2. Metab Brain Dis 30, 1515–1529.

Bushnell, M.C., Ceko, M., and Low, L.A. (2013). Cognitive and emotional control of pain and its disruption in chronic pain. Nat Rev Neurosci 14, 502–511.

Butovsky, O., Jedrychowski, M.P., Moore, C.S., Cialic, R., Lanser, A.J., Gabriely, G., Koeglsperger, T., Dake, B., Wu, P.M., Doykan, C.E., et al. (2014). Identification of a unique TGF-beta-dependent molecular and functional signature in microglia. Nat Neurosci 17, 131–143.

Caldeira, C., Oliveira, A.F., Cunha, C., Vaz, A.R., Falcao, A.S., Fernandes, A., and Brites, D. (2014). Microglia change from a reactive to an age-like phenotype with the time in culture. Front Cell Neurosci 8, 152.

Carey, K.L., Paulus, G.L.C., Wang, L., Balce, D.R., Luo, J.W., Bergman, P., Ferder, I.C., Kong, L., Renaud, N., Singh, S., et al. (2020). TFEB Transcriptional Responses Reveal Negative Feedback by BHLHE40 and BHLHE41. Cell Rep 33, 108371.

Chu, L.W., Cheng, K.I., Chen, J.Y., Cheng, Y.C., Chang, Y.C., Yeh, J.L., Hsu, J.H., Dai, Z.K., and Wu, B.N. (2020). Loganin prevents chronic constriction injury-provoked neuropathic pain by reducing TNF-alpha/IL-1beta-mediated NF-kappaB activation and Schwann cell demyelination. Phytomedicine 67, 153166.

Denk, F., Crow, M., Didangelos, A., Lopes, D.M., and McMahon, S.B. (2016). Persistent Alterations in Microglial Enhancers in a Model of Chronic Pain. Cell Rep 15, 1771–1781.

Doyle, H.H., Eidson, L.N., Sinkiewicz, D.M., and Murphy, A.Z. (2017). Sex Differences in Microglia Activity within the Periaqueductal Gray of the Rat: A Potential Mechanism Driving the Dimorphic Effects of Morphine. J Neurosci 37, 3202–3214.

Echeverry, S., Shi, X.Q., and Zhang, J. (2008). Characterization of cell proliferation in rat spinal cord following peripheral nerve injury and the relationship with neuropathic pain. Pain 135, 37–47.

Filipello, F., Morini, R., Corradini, I., Zerbi, V., Canzi, A., Michalski, B., Erreni, M., Markicevic, M., Starvaggi-Cucuzza, C., Otero, K., et al. (2018). The Microglial Innate Immune Receptor TREM2 Is Required for Synapse Elimination and Normal Brain Connectivity. Immunity 48, 979–991 e978.

Finnerup, N.B., Haroutounian, S., Kamerman, P., Baron, R., Bennett, D.L.H., Bouhassira, D., Cruccu, G., Freeman, R., Hansson, P., Nurmikko, T., et al. (2016). Neuropathic pain: an updated grading system for research and clinical practice. Pain 157, 1599–1606.

Fiore, N.T., and Austin, P.J. (2018). Glial-cytokine-neuronal Adaptations in the Ventral Hippocampus of Rats with Affective Behavioral Changes Following Peripheral Nerve Injury. Neuroscience 390, 119–140.

Fiore, N.T., and Austin, P.J. (2019). Peripheral Nerve Injury Triggers Neuroinflammation in the Medial Prefrontal Cortex and Ventral Hippocampus in a Subgroup of Rats with Coincident Affective Behavioural Changes. Neuroscience 416, 147–167.

Franke, L., el Bannoudi, H., Jansen, D.T., Kok, K., Trynka, G., Diogo, D., Swertz, M., Fransen, K., Knevel, R., Gutierrez-Achury, J., et al. (2016). Association analysis of copy numbers of FC-gamma receptor genes for rheumatoid arthritis and other immune-mediated phenotypes. Eur J Hum Genet 24, 263–270.

Fu, R., Shen, Q., Xu, P., Luo, J.J., and Tang, Y. (2014). Phagocytosis of microglia in the central nervous system diseases. Mol Neurobiol 49, 1422–1434.

Fumagalli, S., Fiordaliso, F., Perego, C., Corbelli, A., Mariani, A., De Paola, M., and De Simoni, M.G. (2019). The phagocytic state of brain myeloid cells after ischemia revealed by superresolution structured illumination microscopy. J Neuroinflammation 16, 9.

Gibson, E.M., Nagaraja, S., Ocampo, A., Tam, L.T., Wood, L.S., Pallegar, P.N., Greene, J.J., Geraghty, A.C., Goldstein, A.K., Ni, L., et al. (2019). Methotrexate Chemotherapy Induces Persistent Tri-glial Dysregulation that Underlies Chemotherapy-Related Cognitive Impairment. Cell 176, 43–55 e13.

Gilchrist, S.E., Goudarzi, S., and Hafizi, S. (2020). Gas6 Inhibits Toll-Like Receptor-Mediated Inflammatory Pathways in Mouse Microglia via Axl and Mer. Front Cell Neurosci 14, 576650.

Goudarzi, S., Gilchrist, S.E., and Hafizi, S. (2020). Gas6 Induces Myelination through Anti-Inflammatory IL-10 and TGF-beta Upregulation in White Matter and Glia. Cells 9.

Grommes, C., Lee, C.Y., Wilkinson, B.L., Jiang, Q., Koenigsknecht-Talboo, J.L., Varnum, B., and Landreth, G.E. (2008). Regulation of microglial phagocytosis and inflammatory gene expression by Gas6 acting on the Axl/Mer family of tyrosine kinases. J Neuroimmune Pharmacol 3, 130–140.

Gu, N., Eyo, U.B., Murugan, M., Peng, J., Matta, S., Dong, H., and Wu, L.J. (2016a). Microglial P2Y12 receptors regulate microglial activation and surveillance during neuropathic pain. Brain Behav Immun 55, 82–92.

Gu, N., Peng, J., Murugan, M., Wang, X., Eyo, U.B., Sun, D., Ren, Y., DiCicco-Bloom, E., Young, W., Dong, H., et al. (2016b). Spinal Microgliosis Due to Resident Microglial Proliferation Is Required for Pain Hypersensitivity after Peripheral Nerve Injury. Cell Rep 16, 605–614.

Guan, Z., Kuhn, J.A., Wang, X., Colquitt, B., Solorzano, C., Vaman, S., Guan, A.K., Evans-Reinsch, Z., Braz, J., Devor, M., et al. (2016). Injured sensory neuron-derived CSF1 induces microglial proliferation and DAP12-dependent pain. Nat Neurosci 19, 94–101.

Gui, W.S., Wei, X., Mai, C.L., Murugan, M., Wu, L.J., Xin, W.J., Zhou, L.J., and Liu, X.G. (2016). Interleukin-1beta overproduction is a common cause for neuropathic pain, memory deficit, and depression following peripheral nerve injury in rodents. Mol Pain 12.

Health, A.I.o., and Welfare (2020). Chronic pain in Australia (Canberra: AIHW).

Hu, L.Y., Zhou, Y., Cui, W.Q., Hu, X.M., Du, L.X., Mi, W.L., Chu, Y.X., Wu, G.C., Wang, Y.Q., and Mao-Ying, Q.L. (2018). Triggering receptor expressed on myeloid cells 2 (TREM2) dependent microglial activation promotes cisplatin-induced peripheral neuropathy in mice. Brain Behav Immun 68, 132–145.

Inoue, K., and Tsuda, M. (2018). Microglia in neuropathic pain: cellular and molecular mechanisms and therapeutic potential. Nat Rev Neurosci 19, 138–152.

Jarjour, N.N., Schwarzkopf, E.A., Bradstreet, T.R., Shchukina, I., Lin, C.C., Huang, S.C., Lai, C.W., Cook, M.E., Taneja, R., Stappenbeck, T.S., et al. (2019). Bhlhe40 mediates tissue-specific control of macrophage proliferation in homeostasis and type 2 immunity. Nat Immunol 20, 687–700.

Jeong, H., Na, Y.J., Lee, K., Kim, Y.H., Lee, Y., Kang, M., Jiang, B.C., Yeom, Y.I., Wu, L.J., Gao, Y.J., et al. (2016). High-resolution transcriptome analysis reveals neuropathic pain gene-expression signatures in spinal microglia after nerve injury. Pain 157, 964–976.

Kehlet, H., Jensen, T.S., and Woolf, C.J. (2006). Persistent postsurgical pain: risk factors and prevention. Lancet 367, 1618–1625.

Keren-Shaul, H., Spinrad, A., Weiner, A., Matcovitch-Natan, O., Dvir-Szternfeld, R., Ulland, T.K., David, E., Baruch, K., Lara-Astaiso, D., Toth, B., et al. (2017). A Unique Microglia Type Associated with Restricting Development of Alzheimer’s Disease. Cell 169, 1276–1290 e1217.

Khoutorsky, A., Sorge, R.E., Prager-Khoutorsky, M., Pawlowski, S.A., Longo, G., Jafarnejad, S.M., Tahmasebi, S., Martin, L.J., Pitcher, M.H., Gkogkas, C.G., et al. (2016). eIF2alpha phosphorylation controls thermal nociception. Proc Natl Acad Sci U S A 113, 11949–11954.

Kinuthia, U.M., Wolf, A., and Langmann, T. (2020). Microglia and Inflammatory Responses in Diabetic Retinopathy. Front Immunol 11, 564077.

Krasemann, S., Madore, C., Cialic, R., Baufeld, C., Calcagno, N., El Fatimy, R., Beckers, L., O’Loughlin, E., Xu, Y., Fanek, Z., et al. (2017). The TREM2-APOE Pathway Drives the Transcriptional Phenotype of Dysfunctional Microglia in Neurodegenerative Diseases. Immunity 47, 566–581 e569.

Lambert, C.S., Philpot, R.M., Engberg, M.E., Johns, B.E., Kim, S.H., and Wecker, L. (2014). Gait analysis and the cumulative gait index (CGI): Translational tools to assess impairments exhibited by rats with olivocerebellar ataxia. Behav Brain Res 274, 334–343.

Lees, J.G., Makker, P.G., Tonkin, R.S., Abdulla, M., Park, S.B., Goldstein, D., and Moalem-Taylor, G. (2017). Immune-mediated processes implicated in chemotherapy-induced peripheral neuropathy. Eur J Cancer 73, 22–29.

Lim, H.Y., Albuquerque, B., Haussler, A., Myrczek, T., Ding, A., and Tegeder, I. (2012). Progranulin contributes to endogenous mechanisms of pain defense after nerve injury in mice. J Cell Mol Med 16, 708–721.

Liu, Y., Zhou, L.J., Wang, J., Li, D., Ren, W.J., Peng, J., Wei, X., Xu, T., Xin, W.J., Pang, R.P., et al. (2017). TNF-alpha Differentially Regulates Synaptic Plasticity in the Hippocampus and Spinal Cord by Microglia-Dependent Mechanisms after Peripheral Nerve Injury. J Neurosci 37, 871–881.

Lopes, D.M., Malek, N., Edye, M., Jager, S.B., McMurray, S., McMahon, S.B., and Denk, F. (2017). Sex differences in peripheral not central immune responses to pain-inducing injury. Sci Rep 7, 16460.

Ma, J., Tanaka, K.F., Shimizu, T., Bernard, C.C., Kakita, A., Takahashi, H., Pfeiffer, S.E., and Ikenaka, K. (2011). Microglial cystatin F expression is a sensitive indicator for ongoing demyelination with concurrent remyelination. J Neurosci Res 89, 639–649.

Ma, Z., Han, Q., Wang, X., Ai, Z., and Zheng, Y. (2016). Galectin-3 Inhibition Is Associated with Neuropathic Pain Attenuation after Peripheral Nerve Injury. PLoS One 11, e0148792.

Makker, P.G., Duffy, S.S., Lees, J.G., Perera, C.J., Tonkin, R.S., Butovsky, O., Park, S.B., Goldstein, D., and Moalem-Taylor, G. (2017). Characterisation of Immune and Neuroinflammatory Changes Associated with Chemotherapy-Induced Peripheral Neuropathy. PLoS One 12, e0170814.

Mannelli, L.D.C., Pacini, A., Bonaccini, L., Zanardelli, M., Mello, T., and Ghelardini, C. (2014). Morphologic features and glial activation in rat oxaliplatin-dependent neuropathic pain. Journal of Pain 14, 1585–1600.

Masuda, T., Iwamoto, S., Yoshinaga, R., Tozaki-Saitoh, H., Nishiyama, A., Mak, T.W., Tamura, T., Tsuda, M., and Inoue, K. (2014). Transcription factor IRF5 drives P2X4R+-reactive microglia gating neuropathic pain. Nat Commun 5, 3771.

Masuda, T., Tsuda, M., Yoshinaga, R., Tozaki-Saitoh, H., Ozato, K., Tamura, T., and Inoue, K. (2012). IRF8 is a critical transcription factor for transforming microglia into a reactive phenotype. Cell Rep 1, 334–340.

Mogil, J.S. (2020). Qualitative sex differences in pain processing: emerging evidence of a biased literature. Nat Rev Neurosci 21, 353–365.

Mols, F., Beijers, T., Vreugdenhil, G., and van de Poll-Franse, L. (2014). Chemotherapy-induced peripheral neuropathy and its association with quality of life: a systematic review. Support Care Cancer 22, 2261–2269.

Monastiriotis, C., Papanas, N., Trypsianis, G., Karanikola, K., Veletza, S., and Maltezos, E. (2013). The epsilon4 allele of the APOE gene is associated with more severe peripheral neuropathy in type 2 diabetic patients. Angiology 64, 451–455.

Monastiriotis, C., Papanas, N., Veletza, S., and Maltezos, E. (2012). APOE gene polymorphisms and diabetic peripheral neuropathy. Arch Med Sci 8, 583–588.

Mor, D., Bembrick, A.L., Austin, P.J., Wyllie, P.M., Creber, N.J., Denyer, G.S., and Keay, K.A. (2010). Anatomically specific patterns of glial activation in the periaqueductal gray of the sub-population of rats showing pain and disability following chronic constriction injury of the sciatic nerve. Neuroscience 166, 1167–1184.

Nishihara, T., Tanaka, J., Sekiya, K., Nishikawa, Y., Abe, N., Hamada, T., Kitamura, S., Ikemune, K., Ochi, S., Choudhury, M.E., et al. (2020). Chronic constriction injury of the sciatic nerve in rats causes different activation modes of microglia between the anterior and posterior horns of the spinal cord. Neurochem Int 134, 104672.

Ochi-ishi, R., Nagata, K., Inoue, T., Tozaki-Saitoh, H., Tsuda, M., and Inoue, K. (2014). Involvement of the chemokine CCL3 and the purinoceptor P2X7 in the spinal cord in paclitaxel-induced mechanical allodynia. Mol Pain 10, 53.

Peng, J., Gu, N., Zhou, L. U B.E., Murugan, M., Gan, W.B., and Wu, L.J. (2016). Microglia and monocytes synergistically promote the transition from acute to chronic pain after nerve injury. Nat Commun 7, 12029.

Plemel, J.R., Stratton, J.A., Michaels, N.J., Rawji, K.S., Zhang, E., Sinha, S., Baaklini, C.S., Dong, Y., Ho, M., Thorburn, K., et al. (2020). Microglia response following acute demyelination is heterogeneous and limits infiltrating macrophage dispersion. Sci Adv 6, eaay6324.

Robinson, C.R., Zhang, H., and Dougherty, P.M. (2014). Astrocytes, but not microglia, are activated in oxaliplatin and bortezomib-induced peripheral neuropathy in the rat. Neuroscience 274, 308–317.

Sorge, R.E., Mapplebeck, J.C., Rosen, S., Beggs, S., Taves, S., Alexander, J.K., Martin, L.J., Austin, J.S., Sotocinal, S.G., Chen, D., et al. (2015). Different immune cells mediate mechanical pain hypersensitivity in male and female mice. Nat Neurosci 18, 1081–1083.

Sousa, C., Golebiewska, A., Poovathingal, S.K., Kaoma, T., Pires-Afonso, Y., Martina, S., Coowar, D., Azuaje, F., Skupin, A., Balling, R., et al. (2018). Single-cell transcriptomics reveals distinct inflammation-induced microglia signatures. EMBO Rep 19.

Staniland, A.A., Clark, A.K., Wodarski, R., Sasso, O., Maione, F., D’Acquisto, F., and Malcangio, M. (2010). Reduced inflammatory and neuropathic pain and decreased spinal microglial response in fractalkine receptor (CX3CR1) knockout mice. J Neurochem 114, 1143–1157.

Sun, L., Wu, Z., Hayashi, Y., Peters, C., Tsuda, M., Inoue, K., and Nakanishi, H. (2012). Microglial cathepsin B contributes to the initiation of peripheral inflammation-induced chronic pain. J Neurosci 32, 11330–11342.

Taves, S., Berta, T., Liu, D.L., Gan, S., Chen, G., Kim, Y.H., Van de Ven, T., Laufer, S., and Ji, R.R. (2016). Spinal inhibition of p38 MAP kinase reduces inflammatory and neuropathic pain in male but not female mice: Sex-dependent microglial signaling in the spinal cord. Brain Behav Immun 55, 70–81.

Taylor, A.M., Mehrabani, S., Liu, S., Taylor, A.J., and Cahill, C.M. (2017). Topography of microglial activation in sensory-and affect-related brain regions in chronic pain. J Neurosci Res 95, 1330–1335.

Tsuda, M., Masuda, T., Kitano, J., Shimoyama, H., Tozaki-Saitoh, H., and Inoue, K. (2009). IFN-gamma receptor signaling mediates spinal microglia activation driving neuropathic pain. Proc Natl Acad Sci U S A 106, 8032–8037.

Tsuda, M., Mizokoshi, A., Shigemoto-Mogami, Y., Koizumi, S., and Inoue, K. (2004). Activation of p38 mitogen-activated protein kinase in spinal hyperactive microglia contributes to pain hypersensitivity following peripheral nerve injury. Glia 45, 89–95.

Tsuda, M., Shigemoto-Mogami, Y., Koizumi, S., Mizokoshi, A., Kohsaka, S., Salter, M.W., and Inoue, K. (2003). P2X4 receptors induced in spinal microglia gate tactile allodynia after nerve injury. Nature 424, 778–783.

Villa, A., Gelosa, P., Castiglioni, L., Cimino, M., Rizzi, N., Pepe, G., Lolli, F., Marcello, E., Sironi, L., Vegeto, E., et al. (2018). Sex-Specific Features of Microglia from Adult Mice. Cell Rep 23, 3501–3511.

Zhang, H., Yoon, M.H., Zhang, H., and Dougherty, P.M. (2012). Evidence that spinal astrocytes but not microglia contribute to the pathogenesis of paclitaxel-induced painful neuropathy. Journal of Pain 13, 293–303.

Zheng, F.Y., Xiao, W.H., and Bennett, G.J. (2011). The response of spinal microglia to chemotherapy-evoked painful peripheral neuropathies is distinct from that evoked by traumatic nerve injuries. Neuroscience 176, 447–454.

