## Supplementary Fig. 1 and 2 for "Sex-specific transcriptome of spinal microglia in neuropathic pain due to peripheral nerve injury"

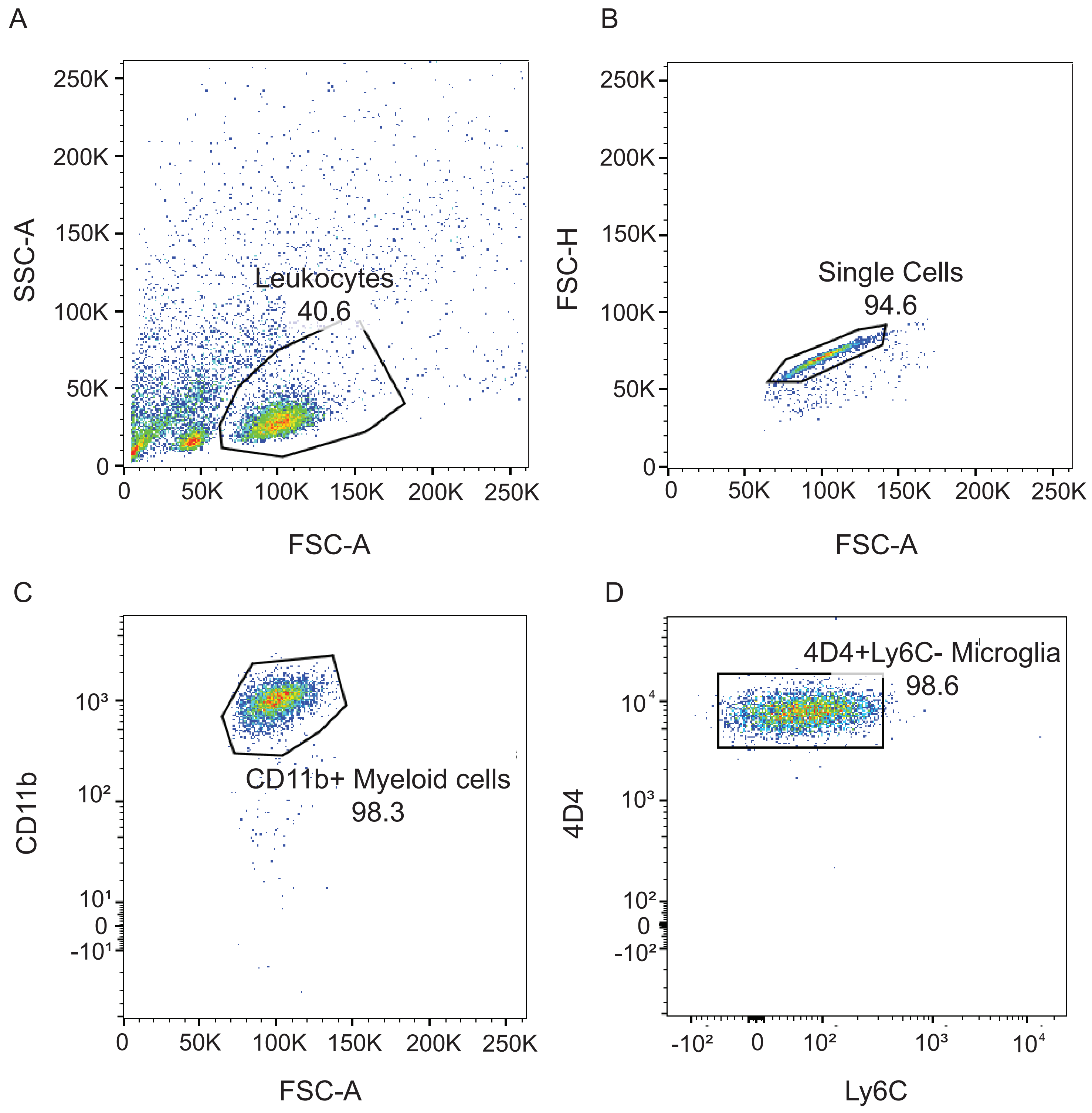
A
B
C
D
SSC-A
FSC-A
FSC-H
Leukocytes
40.6
Single Cells
94.6
CD11b
CD11b+ Myeloid cells
98.3
4D4
Ly6C
4D4+Ly6C- Microglia
98.6
0
50K
100K
150K
200K
250K

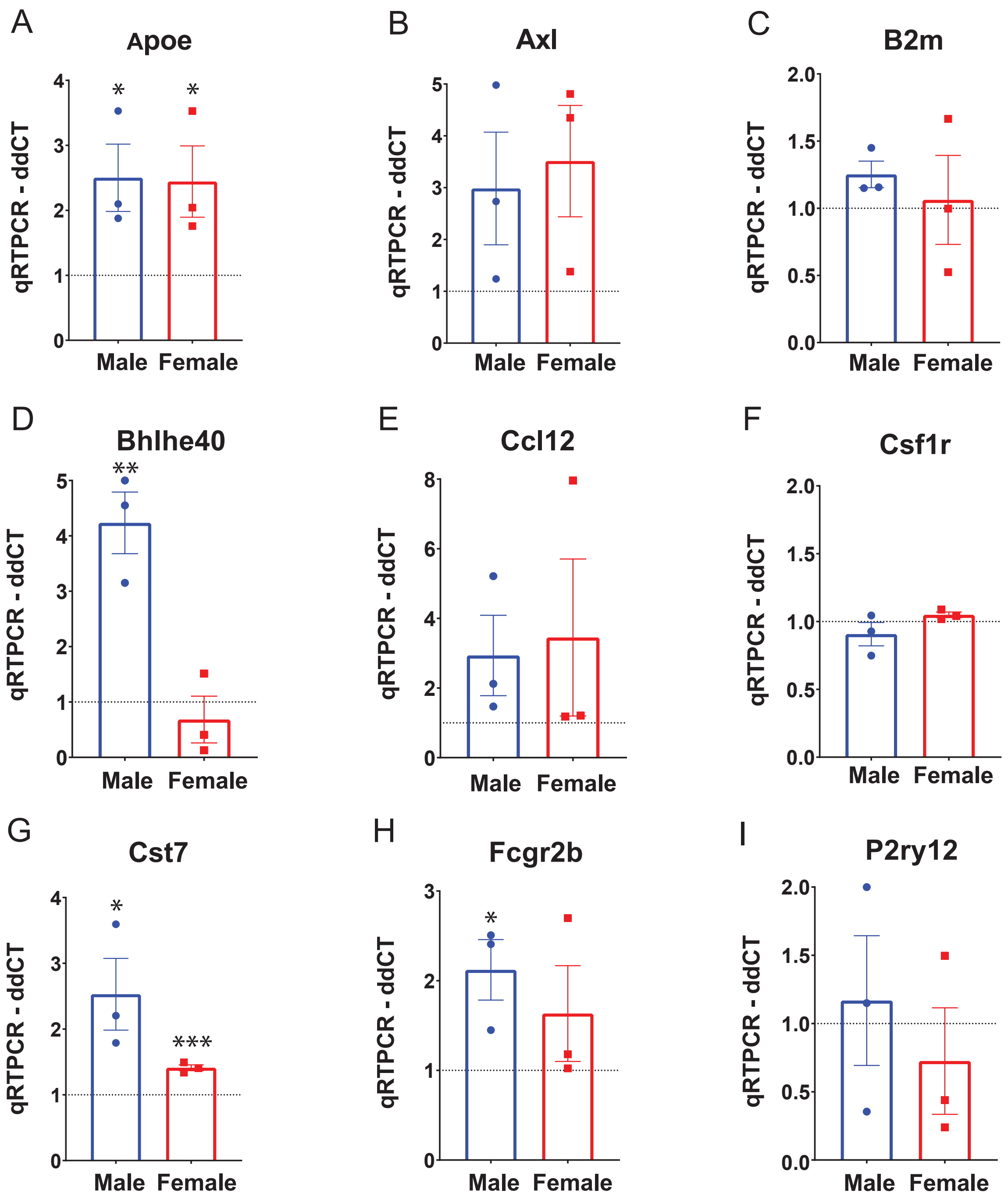

A
Apoe
B
Axl
C
B2m
D
Bhlhe40
E
Ccl12
F
Csf1r
G
Cst7
H
Fcgr2b
I
P2ry12
qRTPCR - ddCT
Male
Female
*
**
***

**Supplementary Figure 1. Microglia isolation with gating strategy.** Microglia were isolated using a percoll gradient and sorted via flow cytometry. (A-D) Representative dot plots illustrating the gating strategy. Markers utilized are (A) cluster of differentiation 11b (Cd11b, myeloid cells), lymphocyte antigen 6 (Ly6C, monocyte/macrophages) and 4D4 (resident microglia).

**Supplementary Figure 2. qRT-PCR validation of gene changes after CCI.** For validation of RNA-seq data, the transcript level of nine selected microglial genes was measured in sorted spinal microglia using qRT-PCR. (A-I) Relative gene expression of microglial genes in the ipsilateral lumbar spinal cord 7 days after CCI in male and female mice relative to *Gapdh* expression and sham mice (as indicated by the dotted line at y = 1). qRCR confirmed an increase in *Apoe*, *Bhlhe40*, *Cst7* and *Fcgr2b* in male mice and in *Apoe* and *Cst7* in female mice after CCI. Data are presented as mean ddCT ± SEM (* $p < 0.05$, ** $p < 0.01$ and *** $p < 0.001$).
